# Proteolytic processing of Argonautes in multiphase condensates promotes transgenerational epigenetic inheritance

**DOI:** 10.64898/2026.01.01.697165

**Authors:** Boyuan Deng, Xufang Niu, Haiqian Ma, Lufan Shi, Changfeng Zhao, Wanlin Chen, Xinru Li, Yuntao Liang, Shuyu Hu, Ken Chen, Yonglin Liang, Yanhong Qiu, Haoxiang Jia, Gang Wan

**Affiliations:** Guangdong Province Key Laboratory of Pharmaceutical Functional Genes, MOE Key Laboratory of Gene Function and Regulation, State Key Laboratory of Biocontrol, School of Life Sciences, Sun Yat-sen University, GuangZhou, GuangDong 510275, China; Innovation Center for Evolutionary Synthetic Biology, School of Life Sciences, Sun Yat-sen University, Guangzhou, China

## Abstract

Epigenetic information can be memorized in perinuclear condenates to transmit across generations to modulate gene expression in descenent. Here, we identify a conserved N terminal proteolytic mechanism in *Caenorhabditis elegans* that is essential for small RNA-mediated transgenerational epigenetic inheritance (TEI) and for maintaining germline immortality. Two conserved, perinuclear germ granule subcompartments-enriched N-terminal processing peptidases, APP-1 and DPF-3, act cooperatively to trim N-terminal intrinsically disordered regions of the Argonaute proteins WAGO-1, WAGO-3, and WAGO-4. This processing shields these Argonautes from proteasome-mediated protein degradation, prevents the loss of their associated small RNAs, and promotes their accumulation in germ granules subcompartments. In contrast, unprocessed WAGOs are recognized and destabilized by the conserved E3 ubiquitin ligases RNF-1 and UBR-7. We propose that this cooperative proteolytic mechanism adds another layer of sspatiotemporal regulation over small RNA-directed epigenetic inheritance and may represents a broadly conserved strategy for regulating protein homeostasis and gene expression programs.

## Introduction

Genetic information is classically transmitted from parents to offspring through DNA in a mendelian manner. However, increasing evidence indicates that epigenetic information-such as DNA and histone modifications, as well as small RNAs including small interfering RNAs (siRNAs), microRNAs, and tsRNAs-can also be inherited across generations to regulate gene expression independently of DNA sequence^1–4^. Such non-mendelian epigenetic inheritance can be triggered by environmental stimuli, potentially enabling offspring to adapt to changing conditions and having implications in human health and evolution^5–8^. The persistence of epigenetic inheritance varies, ranging from one generation to several; when inheritance extends beyond three generations, it is typically referred to as transgenerational epigenetic inheritance (TEI)^1,2,9^. Despite growing examples of TEI, the molecular mechanisms that establish, maintain and regulate it remain largely elusive.

One of the most prominent examples of TEI is RNA interference (RNAi) inheritance in *Caenorhabditis elegans* (*C. elegans*)^10–12^. RNAi is a conserved, small RNA-mediated, post-transcriptional gene silencing mechanism^13^. In *C. elegans*, siRNAs known as 22G-siRNAs are generated from environmental double-stranded RNAs (dsRNAs) or endogenous PIWI-interacting RNAs (piRNAs) through RNA-dependent RNA polymerase (RdRP)-mediated amplification^14–16^. A class of Argonaute proteins, the WAGOs, bind these 22G-siRNAs and guide them to complementary target RNAs via Watson-Crick base pairing to initiate silencing^17^. These silencing signals can be transmitted to progeny and perpetuate gene repression across generations through WAGOs-containing machineries that operate both in the nucleus and cytoplasm. In the nucleus, the Argonaute HRDE-1 associates with NRDE-1, NRDE-2, and NRDE-4 to inhibit nascent transcription at the elongation stage, promoting deposition of heterochromatic marks H3K9me3 and H3K27me3^18–22^. In the cytoplasm, germ granules-membraneless organelles or biomolecular condensate formed through liquid-liquid phase separation-serve as hubs for RNAi triggered gene silencing and its memory^23,24^. Moreover, disruption of genes required for TEI leads to progressive germline sterility, particularly under elevated temperatures^18,25–27^, underscoring the essential role of RNAi inheritance in maintaining germline immortality.

Germ granules are conserved, typically perinuclear condensates present in most, if not all, animals and are thought to surveil and process nascent mRNAs as they exit the nuclear pore^28,29^. In *C. elegans*, germ granules are multiphase condensates composed of at least seven functionally distinct yet interconnected subcompartments, including P granules^30^, Z granule^26^, Mutator foci^31^, SIMR foci^32^, D granule^33,34^, E granule^35^, and P body^36^. Each subcompartment hosts specialized small RNA pathways that coordinate the initiation, amplification, inheritance, and regulation of gene-silencing signals^25,37^. For example, WAGO-4 and the conserved RNA helicase ZNFX-1 maintain silencing memory within Z granules^26,38–40^, whereas WAGO-1 and WAGO-3 localize to P granules and also contribute to RNAi inheritance^41,42^. Mutator foci, marked by MUT-16, serve as centers for small RNA amplification^31^. The multiphase organization of germ granules is proposed to coordinate complex and potentially competing small RNA pathways^25,26^. Each subcompartment contains dozens to hundreds of protein components, as revealed by proximity-dependent biotin labeling and immunoprecipitation–coupled mass spectrometry^25,37,43–46^. However, the functions of most of these components remain unknown, and the biological significance of the multiphase organization of germ granules is still poorly understood.

In addition to serving as hubs for small RNA pathways, germ granules also concentrate proteins that modulate small RNA machinery activity and maintain granule homeostasis^25,47,48^. One such protein is the conserved dipeptidyl peptidase DPF-3, a P-granule-localized member of the dipeptidyl peptidase IV serine protease family^48^. The N termini of proteins are frequently subjected to co- or post-translational modifications, such as N-terminal acetylation^49^, or can function as N-degrons that determine protein stability^50^. These N-terminal residues are crucial for protein localization, protein-protein interactions, and protein degradation^50^. In addition, the N-terminal amino acids often undergo peptidase processing; for instance, co-translational removal of the initiating methionine by methionine aminopeptidase is a common maturation step for nascent polypeptides^51^.

DPF-3 specifically removes dipeptides (Xaa-Ala or Xaa-Pro, where Xaa represents any amino acid) from protein N termini. Loss of *dpf-3* causes reduced fertility and elevated transposon activity in *C. elegans*^48^. WAGO-1 and WAGO-3 have been identified as DPF-3 substrates; DPF-3-mediated processing influences WAGO-1 siRNA loading and WAGO-3 protein stability, although the underlying mechanisms remain unclear^48^. Another conserved N-terminal peptidase, APP-1, can cleave Xaa-Pro bonds at the N terminus^52^ and may therefore target a similar set of substrates as DPF-3, potentially acting in a competitive or cooperative manner. Proteomic analysis of germ granules has identified APP-1, suggesting that it may be a component of these condensates^25,53^. Loss of APP-1 leads to replication-associated genome instability-a phenotype conserved in both *C. elegans* and human cells-and results in fertility defects^54^. However, specific APP-1 substrates in either *C. elegans* or human have not yet been identified. The roles of these N-terminal processing enzymes, and whether they act cooperatively to regulate germline proteostasis and small RNA inheritance, remain unexplored.

Here, we show that germ granules-enriched APP-1 and DPF-3 act cooperatively to promote small RNA-based TEI and maintain germline immortality in *C. elegans*. We identify three germ granules-enriched Argonaute proteins-WAGO-1, WAGO-3, and WAGO-4-as shared substrates of APP-1 and DPF-3. Loss of APP-1- or DPF-3-mediated processing triggers the degradation of WAGO-1, WAGO-3, and WAGO-4 through E3 ligases-RNF-1 and UBR-7-dependent proteasomal pathways, driven by their N-terminal proline-rich intrinsically disordered regions. Together, our findings uncover a conserved cooperative proteolytic mechanism that safeguards Argonautes and associated small stability in the condensate to ensure epigenetic memory.

## Results

### Two conserved N-terminal processing peptidases are critical for RNAi inheritance

Germ cells are immortal cell lineage, and germline immortality is under genetic control^55^. Previous studies have shown that several genes encoding germ granule components are essential for promoting germline immortality in *C. elegans*^25–27,32,45,47,56–58^. To further investigate the role of germ granule components, we previously employed TurboID-based proximity biotin labeling to identify protein constituents within each PZM germ granule subcompartments^25^. Deletion mutants were generated for genes encoding the most enriched germ granule components, and these mutants were subjected to germline immortality assays^25^. In these assays, we screened for mutants exhibiting progressive fertility decline across generations, culminating in sterility after three to ten generations at 25°C. Out of 50 fertile mutants, 8 were identified as essential for germline immortality^25^. To expand our understanding of germ granule functions, we generated additional mutants targeting other germ granule components and subjected them to similar germline immortality assays. One of the identified genes encodes the conserved N-terminal aminopeptidase APP-1^54^. When *app-1(-)* animals were maintained at 25°C, their brood size progressively declined with each generation, ultimately resulting in sterility after three to six generations (Figures 1A and 1C), suggesting that APP-1 is critical for maintaining germline immortality. A previous study reported that loss of APP-1 results in increased DNA damage and genome instability^54^. To determine whether the loss of germline immortality in *app-1(-)* animals stems from DNA damage or dysregulated epigenetic inheritance, we conducted a temperature-shift experiment. Specifically, *app-1(-)* animals were reared at 25°C for three generations (F3), approaching sterility, and then transferred to 20°C (Figure 1D). Notably, their brood size progressively increased with each subsequent generation at 20°C (Figure 1D). This cycle was repeated by shifting the animals back to 25°C and then to 20°C again. Brood size consistently decreased at 25°C and increased at 20°C (Figure 1D). These results indicate that APP-1 supports germline immortality by regulating epigenetic inheritance rather than solely mitigating DNA damage.

**Figure 1.**
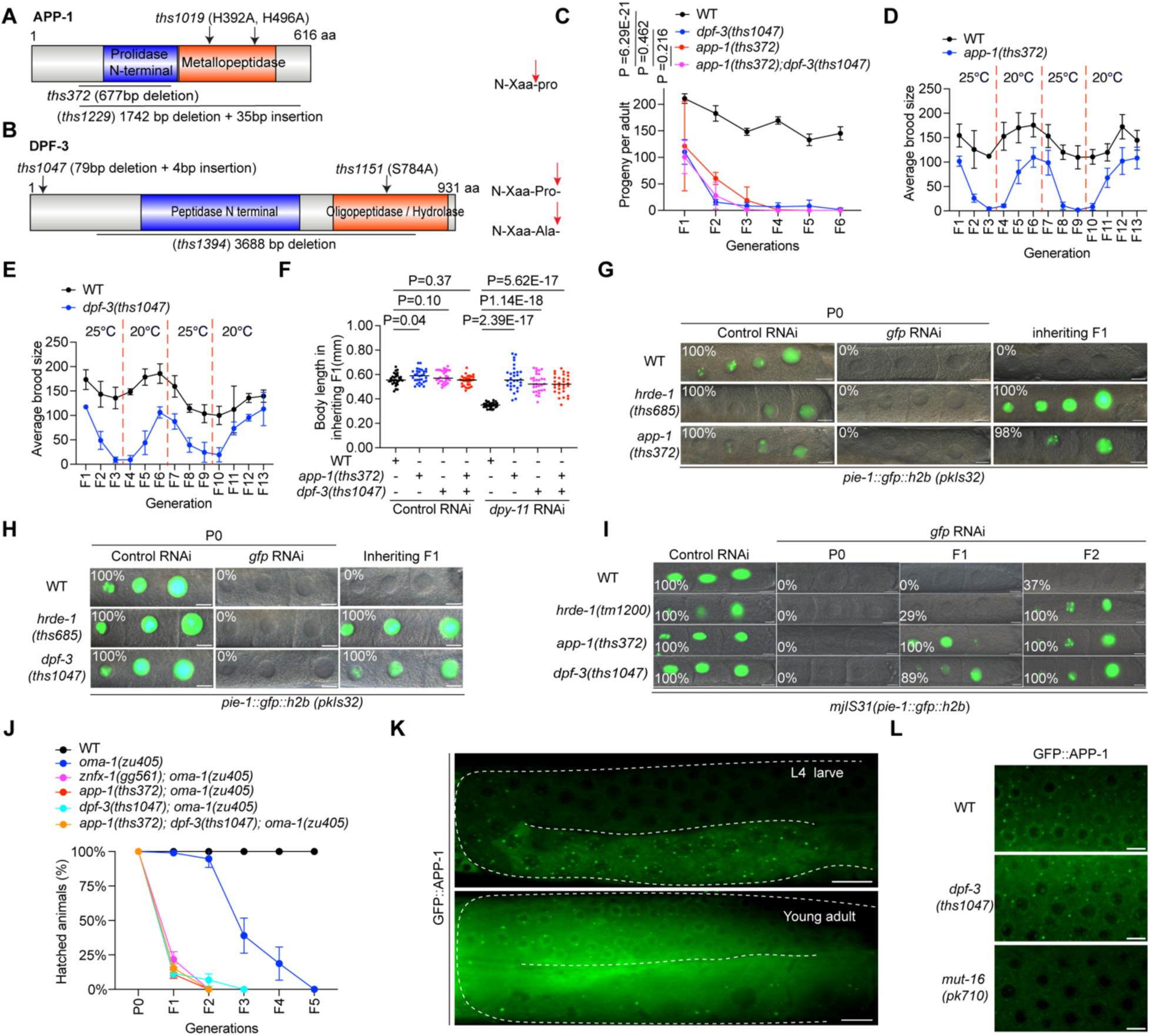
Two conserved N-terminal peptidases promote small RNA-based epigenetic inheritance and germline immortality. (A-B) Schematic diagram of APP-1 and DPF-3 domains, with alleles used in this study indicated. Red arrows indicate predicted cleavage sites of APP-1 and DPF-3. (C) Brood sizes of WT, *app-1(-)*, *dpf-3(-)*, and *app-1(-)*; *dpf-3(-)* animals grown at 25 °C across generations. n = 3 biological replicates. P values were determined by one-way ANOVA. (D-E) Brood sizes of WT and *app-1(-)* animals (D), or WT and *dpf-3(-)* animals (E), maintained at 25 °C or 20 °C for the indicated generations. n = 3 biological replicates. (F) Animals were subjected to *dpy-11* RNAi, progeny were grown on non-RNAi plates. Body length of F1 animals was quantified at L4 larval stage using ImageJ (30 animals per genotype). Student’s t test, two-tailed. (G-H) WT, *hrde-1(-)*, and *app-1(-)* animals (G) or WT, *hrde-1(-)*, and *dpf-3(-)* animals (H) expressing *gfp::h2b* in the germline (*pkIS32*) were exposed to *gfp* RNAi; progeny were maintained on no RNAi plates. The percentage of animals expressing GFP in −1 to −3 oocytes were scored (50 animals per replicate, n = 3 biological replicates). Scale bar, 10μm. (I) WT, *hrde-1(-)*, *app-1(-)*, and *dpf-3(-)* animals expressing *gfp::h2b* in the germline (*mjIS31*) were treated as in (G-H). GFP expression in −1 to −3 oocytes was scored (50 animals per replicate, n = 3 biological replicates). Scale bar, 10μm. (J) Animals carrying *oma-1(zu405)* were treated with *oma-1* dsRNA and maintained at 20°C. The percentage of hatched progeny were scored across generations (n = 3 biological replicates). (K) Fluorescence imaging of L4 and young adult animals expressing GFP::APP-1. n > 3 animals. Scale bar, 10μm. (L) Fluorescence imaging of young adult WT, *dpf-3(-)*, and *mut-16(-)* animals expressing GFP::APP-1. n > 3 animals. Scale bar, 5μm.

APP-1 is a conserved N-terminal aminopeptidase that specifically cleaves peptide bonds between an amino acid (denoted as Xaa, one of the 20 standard amino acids) and proline at the N-terminus (Figure 1A)^52,54^. Similarly, DPF-3 is another N-terminal peptidase localized to germ granule subcompartments (P granules) and catalyzes the cleavage of two-amino-acid peptides, either Xaa-Pro or Xaa-Ala, from the N-terminus^48^. Given the similarity in substrate specificity, APP-1 and DPF-3 may process overlapping sets of substrate. This prompted us to investigate whether DPF-3 plays a similar role in promoting germline immortality. To address this, we generated a mutant for *dpf-3* that was presumably null (Figure 1B). Similar to *app-1(-)* mutants, the progeny of *dpf-3(-)* animals displayed progressive declines in brood size over generations, culminating in sterility after three to six generations at 25°C (Figure 1C). Loss of both *app-1* and *dpf-3* also resulted in germline immortality defects, but no statistically significant difference was observed between the double mutants (*app-1(-)*; *dpf-3(-)*) and the single mutants (*app-1(-)* or *dpf-3(-)*) (Figure 1C). Given that the loss of DPF-3 also causes DNA damage^48^, we investigated whether the germline immortality defects observed in *dpf-3(-)* animals occurred at an epigenetic level. Temperature shift experiments revealed that the disruption of germline immortality was indeed epigenetically regulated (Figure 1E). In summary, we identified two conserved N-terminal processing peptidases, APP-1 and DPF-3, that function, at least partially redundantly, to promote germline immortality in *C. elegans*.

Given that several genes promoting germline immortality are also required for RNAi-mediated epigenetic inheritance^18,47,53,59^, we asked whether APP-1 and DPF-3 play a role in this process. Four lines of evidence support the involvement of APP-1 and DPF-3 in promoting RNAi-based epigenetic inheritance. First, *dpy-11* encodes a hypodermal protein that regulates body length^60^. Wild-type (WT) animals exposed to *dpy-11* RNAi exhibit shorter body length compared to those treated with control RNAi, and this silencing effect is heritable in the F1 generation (Figure 1F)^61^. When *app-1(-)* or *dpf-3(-)* animals were subjected to *dpy-11* RNAi, they displayed the Dumpy (Dpy) phenotype, similar to WT animals (Figure 1F). However, while WT animals remained Dpy in the inheriting F1 generation, *app-1(-)* and *dpf-3(-)* animals had a longer body length compared to WT (Figure 1F), suggesting *app-1* or *dpf-3* are required for *dpy-11* RNAi inheritance. Second, *gfp* RNAi targeting a germline expressed *gfp::h2b* transgene (*pkIS32*) induces GFP silencing, which is heritable for ~5 generations, thus serving as a reporter for TEI^18^. TEI was abolished upon depletion of the nuclear Argonaute HRDE-1(Figures 1G-H)^18^ and was similarly lost in *app-1(-)* or *dpf-3(-)* mutants (Figures 1G-H). Furthermore, APP-1 and DPF-3 were required for *gfp* RNAi-mediated TEI in a second single-copy *gfp::h2b* transgene (*mjIS31*) (Figure 1I)^14,26^. These results suggest that APP-1 and DPF-3 promote TEI. Third, *oma-1(zu405)* encodes a temperature sensitive dominant OMA-1 mutant proteins that resist degradation, causing embryonic arrest at restrictive temperatures (20°C)^62^. *oma-1* RNAi silences the mutant OMA-1 protein, allowing animals exposed to *oma-1* RNAi and their progeny to grow at 20°C. WT animals maintained this silencing and continued to grow at 20°C for ~5 generations, consistent with previous reports (Figure 1J)^12,26^. In contrast, while embryos from *app-1(-)* or *dpf-3(-)* animals treated with *oma-1* RNAi hatched normally, embryos from the progeny of these treated mutants arrested after two generations at 20°C (Figure 1J). This indicates that APP-1 and DPF-3 are required for RNAi inheritance targeting endogenous gene *oma-1*. Finally, piRNAs can induce multigenerational epigenetic inheritance by templating the synthesis of siRNAs through RDRP^14,16^. This process can be monitored using a *gfp::h2b* piRNA sensor, which is typically silenced. GFP silencing was reversed in *hrde-1(-)*, *app-1(-)*, *dpf-3(-)*, and *app-1(-); dpf-3(-)* animals, although the desilencing was less pronounced in *app-1(-)*, *dpf-3(-)*, and *app-1(-); dpf-3(-)* animals compared to *hrde-1(-)* animals (Figures S1A-C). Taken together, these results suggest that APP-1 and DPF-3 are essential for small RNA-based epigenetic inheritance and germline immortality.

### APP-1 and DPF-3 peptidase activity is essential for RNAi inheritance

Peptidases like DPF-3 can function independently of their enzymatic activity^63^. To determine whether the peptidase enzyme activity of APP-1 and DPF-3 is required for epigenetic inheritance and germline immortality, we introduced point mutations to key residues predicted to be critical for their enzymatic activity using CRISPR/Cas9 (Figure 1A)^48,54^. Specifically, we generated *app-1(H392A, H496A)* mutants (with Histidine 392 and Histidine 496 replaced by Alanine) and *dpf-3(S784A)* mutants (with Serine 784 replaced by Alanine), which have been shown to abolish APP-1 and DPF-3 peptidase activity, respectively^48,54^. When *app-1(H392A, H496A)* or *dpf-3(S784A)* animals were grown at 25°C, they progressively lost fertility, becoming completely sterile by the sixth generation, similar to *app-1* and *dpf-3* deletion mutants (Figures S1D-E). Similarly, *app-1(H392A, H496A)* and *dpf-3(S784A)* mutants were defective in *dpy-11* RNAi inheritance (Figures S1F-G). When these mutants expressing *gfp::h2b* were subjected to *gfp* RNAi, the resulting gene silencing failed to be inherited (Figures S1H-I). Finally, *app-1(H392A, H496A)*; *oma-1(zu405)* and *dpf-3(S784A)*; *oma-1(zu405)* animals exposed to *oma-1* RNAi were able to develop embryos; however, embryos in subsequent generations arrested after two generations without *oma-1* RNAi treatment (Figure S1J). This phenotype was similar to that observed in *app-1(-)* and *dpf-3(-)* animals, indicating that the peptidase enzyme activity of APP-1 and DPF-3 is essential for TEI. In conclusion, these results suggest that the peptidase activity of APP-1 and DPF-3 is critical for small RNA-based epigenetic inheritance and the maintenance of germline immortality.

### APP-1 is enriched in Mutator foci

While a previous study reported that DPF-3 localizes to the cytoplasm and is enriched in P granules^48^, the localization of APP-1 remained unknown. To determine the organismal and subcellular localization of APP-1, we introduced a GFP tag immediately downstream of the endogenous *app-1* start codon and generated animals expressing GFP::APP-1 using CRISPR/Cas9. The resultant GFP::APP-1 fusion protein was functional, as *gfp::app-1* animals behaved like WT animals in the *dpy-11* RNAi inheritance assay (Figure S3A). Fluorescence imaging using spinning disc microscopy revealed that GFP::APP-1 was expressed in the germline and during embryogenesis (Figures 1K and S2A). In the germline, GFP::APP-1 displayed diffuse cytoplasmic expression, with a subset of the protein enriched in perinuclear condensates (Figure 1K). During embryogenesis, GFP::APP-1 was diffusely expressed in the cytoplasm of both somatic and germline blastomeres, with no asymmetrical distribution observed (Figure S2A). GFP::APP-1 expression was also observed in various somatic tissues, which was diffusely localized in the cytoplasm (Figure S2A).

The germ granules in *C. elegans* germline are composed of at least four distinct perinuclear sub-compartments: P granules^26^, Z granules^26^, Mutator foci^31^ and SIMR foci^32^. To determine which sub-compartment APP-1 concentrate to, we generated animals expressing GFP::APP-1 and marker proteins for each sub-compartment using genetic cross. Fluorescence imaging showed that GFP::APP-1 localized adjacent to PGL-1, ZNFX-1 and SIMR-1, markers of P granule, Z granule and SIMR foci, respectively (Figure S2B). However, GFP::APP-1 co-localized with MUT-2::mCherry, a marker of Mutator foci (Figure S2B). ImageJ-based co-localization analysis confirmed that the perinuclear GFP::APP-1 fluorescence signal overlapped with MUT-2::mCherry fluorescence signal (Figure S2C). MUT-16 is a scaffold protein essential for the integrity of Mutator foci^31^. We found that the perinuclear GFP::APP-1 granular signal was disrupted in *mut-16(-)* animals but not in *dpf-3(-)* animals (Figure 1L). In summary, these results suggest that APP-1 is diffusely expressed in the cytoplasm of the germline, with a portion specifically enriched in Mutator foci.

### APP-1 and DPF-3 are required for the transgenerational transmission or maintenance of siRNAs

During TEI, siRNAs targeting silenced genes can persist for multiple generations^17^. To investigate whether APP-1 and DPF-3 regulate siRNAs production or transmission during TEI, we performed small RNA sequencing on WT, *app-1(-)*, and *dpf-3(-)* animals exposed to *oma-1* dsRNA. In the directly exposed P0 generation, levels of *oma-1*-targeting siRNAs were comparable-or modestly elevated-in *app-1(-)* and *dpf-3(-)* mutants relative to WT (Figure 2A), suggesting that siRNA biogenesis in response to dsRNA remains intact in the absence of APP-1 or DPF-3. However, in the F1 progeny that inherited the silencing signal but were not directly exposed to dsRNA, *oma-1*-targeting siRNAs were significantly depleted-by approximately 95% in *app-1(-)* and *dpf-3(-)* mutants compared to WT controls (Figure 2A). These findings indicate that while APP-1 and DPF-3 are dispensable for initial siRNA production, they are essential for the transmission or maintenance of siRNAs across generations.

**Figure 2.**
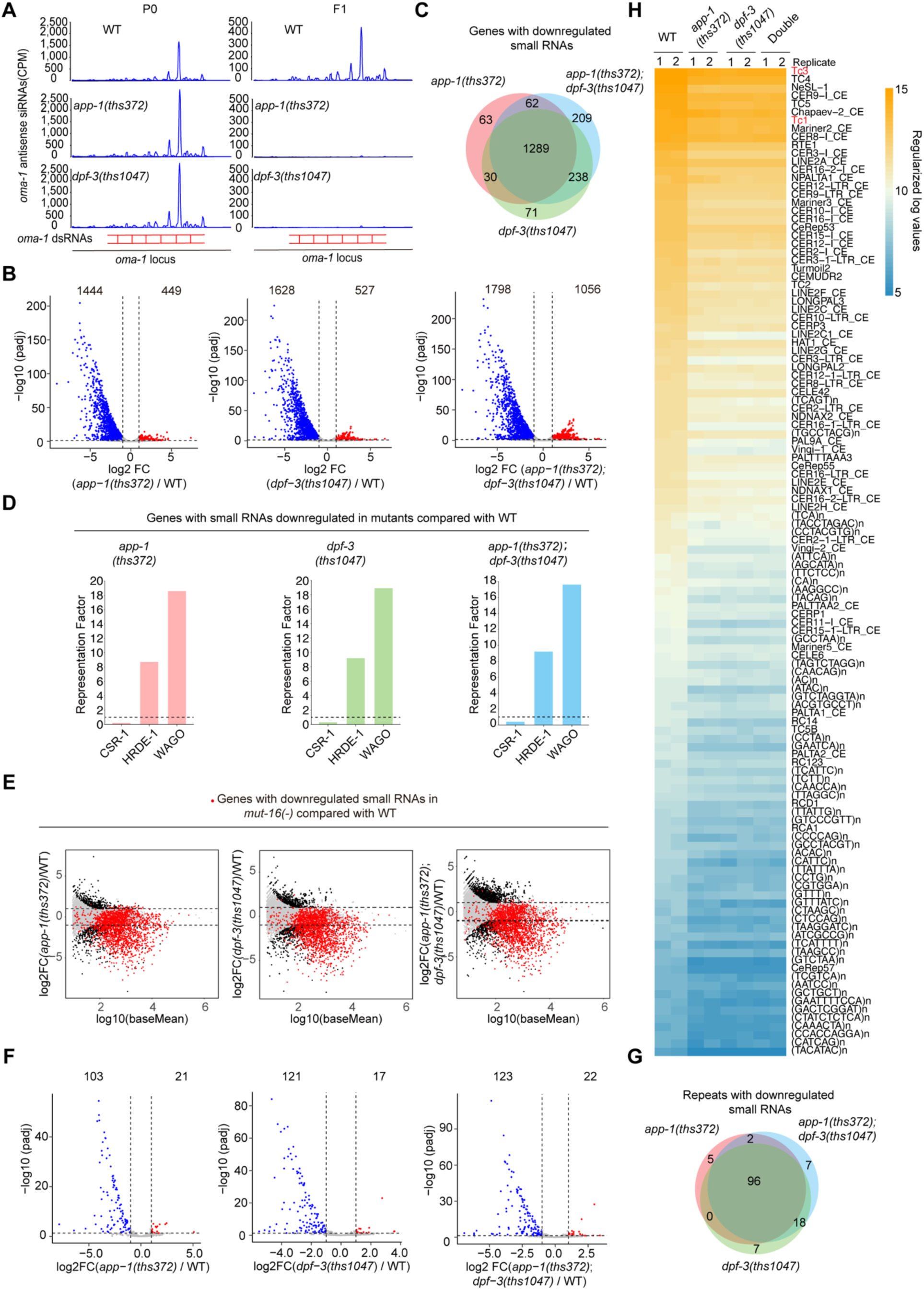
Small RNA-seq analysis of APP-1 and DPF-3 regulated small RNAs. (A) WT, *app-1(-)*, and *dpf-3(-)* animals were subjected to *oma-1* RNAi, antisense *oma-1* small RNAs mapped to the *oma-1* locus are shown for the P0 and inheriting F1 generations. (B) Volcano plot showing protein-coding genes differentially regulated by small RNAs in *app-1(-)*, *dpf-3(-)*, and *app-1(-)*; *dpf-3(-)* animals compared with WT. Dotted line indicate log2 Fold change >= 1 or <= −1. (C) Venn diagram showing protein-coding genes with reduced small RNA levels in *app-1(-)*, *dpf-3(-)*, and *app-1(-)*; *dpf-3(-)* animals relative to WT. (D) Representative factor analysis comparing genes with reduced small RNAs in *app-1(-)*, *dpf-3(-)*, and *app-1(-)*; *dpf-3(-)* animals relative to WT with genes regulated by CSR-1 class-, HRDE-1 class-, and WAGO class-small RNAs. (E) Dot plot showing protein-coding genes with differential small RNA levels in *app-1(-)*, *dpf-3(-)*, and *app-1(-)*; *dpf-3(-)* relative to WT and genes with differential small RNAs levels in *mut-16(-)* relative to WT. Dotted lines indicate log2 Fold change >= 1 or <= −1. (F) Volcano plot showing transposons and repetitive sequences differentially regulated by small RNAs in *app-1(-)*, *dpf-3(-)* and *app-1(-)*; *dpf-3(-)* animals compared with WT. Dotted line indicate log2 Fold change >= 1 or <= −1. (G) Venn diagram showing transposons and repetitive sequences with reduced small RNA levels in *app-1(-)*, *dpf-3(-)*, and *app-1(-)*; *dpf-3(-)* animals relative to WT. (H) Heatmap of transposons and repetitive sequences that exhibit reduced small RNA abundance in *app-1(-)*, *dpf-3(-)* and *app-1(-)*; *dpf-3(-)* animals compared with WT.

*C. elegans* expresses abundant endogenous small RNAs that associate with WAGOs to mediate co-transcriptional or post-transcriptional gene silencing^13^. Many factors involved in small RNA-mediated gene regulation are also linked to TEI and germline immortality^18,26,27,39^. A previous study identified a subset of endogenous small RNAs regulated by DPF-3 at the L4 larval stage at 15 °C^48^; however, whether APP-1 similarly influences endogenous small RNA populations remained unclear. To address this, we performed small RNA-seq on total RNA from WT, *app-1(-)*, *dpf-3(-)*, *app-1(-);dpf-3(-)* young adult animals reared at 20°C. Mapping these small RNAs to gene loci revealed 1,893, 2,155, and 2,854 genes that were differentially targeted in *app-1(-)*, *dpf-3(-)*, *app-1(-);dpf-3(-)* animals respectively, compared to WT (log2FoldChange ≥ 1 or ≤ −1, adjusted P value < 0.05) (Figure 2B). Specifically, we identified 1,444, 1,628, and 1,798 genes exhibiting reduced small RNA levels and 449, 527, and 1056 genes showing increased levels in *app-1(-)*, *dpf-3(-)*, *app-1(-);dpf-3(-)* animals, respectively-revealing an ~1.7-3.2 fold bias toward downregulation. A majority (1289, 72%-89%) of the downregulated targets overlapped among all three mutants (Figure 2C). To further define the role of APP-1 and DPF-3, we compared their regulated gene sets with previously characterized targets of endogenous RNAi factors, including CSR-1^64^, HRDE-1^18^, and WAGOs^65^. Genes with reduced small RNA levels in *app-1(-)*, *dpf-3(-)*, and *app-1(-)*; *dpf-3(-)* mutants were significantly enriched for HRDE-1 and WAGO targets, but not CSR-1 targets (Figure 2D). Together, these results suggest that APP-1- and DPF-3-mediated N-terminal processing promotes the stability of a subset of HRDE-1- and WAGO-associated small RNAs.

Loss of APP-1 in *C. elegans* and human cells leads to genome instability; however, the underlying mechanism remains unclear^54^. Similarly, DPF-3 in *C. elegans* protects genome integrity^48^. Small RNA-based gene silencing plays a crucial role in controlling selfish genetic elements, such as transposons and repetitive sequences, which, if left unchecked, can cause genome instability^65^. Notably, the small RNA machinery, concentrated in Mutator foci, has been linked to the silencing of transposons and repetitive sequences^31,66^. We investigated whether small RNAs regulated by APP-1 and DPF-3 overlapped with those regulated by MUT-16^32^. Indeed, genes with small RNAs downregulated in *app-1(-)*, *dpf-3(-)*, *app-1(-); dpf-3(-)* mutants were significantly enriched for MUT-16-regulated genes (Figure 2E). Additionally, we observed a reduction in small RNAs targeting specific transposons (such as TC1 and TC3) and repetitive sequences in *app-1(-)*, *dpf-3(-)*, *app-1(-); dpf-3(-)* animals compared to WT animals (Figures 2F-H). These findings suggest that APP-1- and DPF-3-regulated siRNAs may contribute, at least in part, to maintaining genome stability.

### Defective processing of WAGO-1/3/4 phenocopies APP-1 and DPF-3 loss in RNAi inheritance and germline immortality

Given that methionine can be co-translationally removed^51^, proteins with N-terminal sequences such as MP, MXaaP, MA, or MXaaA may serve as potential DPF-3 substrates, while those with MP or MXaaP at the N-terminus may be substrates of APP-1. Furthermore, DPF-3 cleavage could expose Xaa-Pro, converting previously non-substrate proteins of APP-1 into APP-1 substrates, and vice versa. To identify substrates involved in TEI and germline immortality, we analyzed proteins with such N-terminal motifs. Among the candidates that may contribute to these processes, we identified WAGO-1^41,67^, WAGO-3^42^, WAGO-4^38^, SET-25^68^, and PID-5^53^. To confirm whether these proteins are APP-1 or DPF-3 substrates, we introduced point mutations at predicted processing sites, replacing proline or alanine residues with glycine or glutamine (P to G, A to E) respectively (Figure 3A). The mutations were predicted to abolish processing by APP-1 and DPF-3. We evaluated the resulting mutants using the *dpy-11* RNAi inheritance assay. Notably, *wago-1(P3G)*, *wago-3(P2G, A3E)*, and *wago-4(P2G, A3E)* mutants-but not other processing-defective mutants-exhibited defects or modest defects in *dpy-11* RNAi inheritance (Figures S4A-D). These results indicate that WAGO-1, WAGO-3, and WAGO-4 are likely APP-1 or DPF-3 substrates involved in RNAi inheritance.

**Figure 3.**
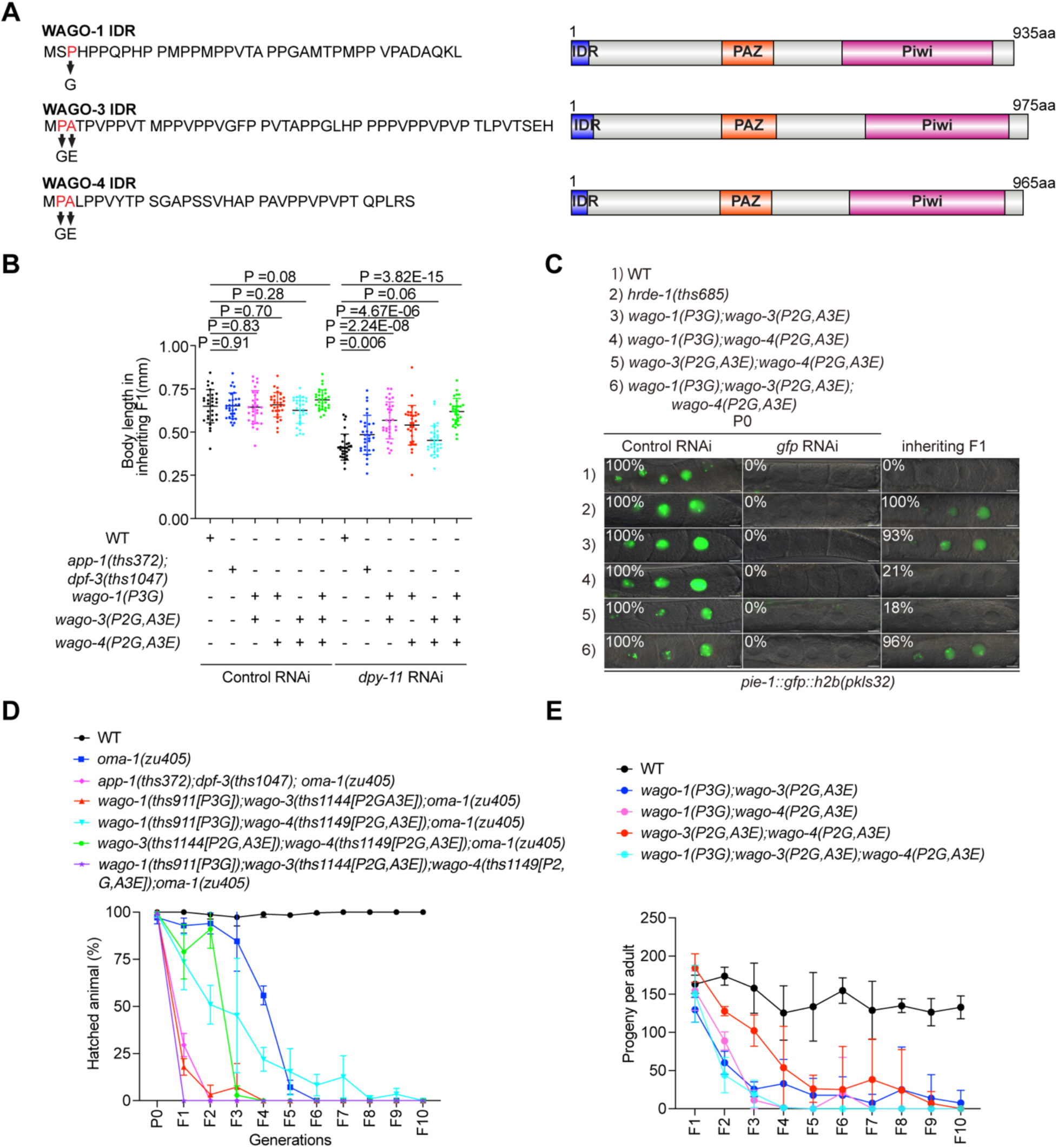
Impaired WAGO-1/3/4 processing phenocopies APP-1 and DPF-3 loss in RNAi inheritance and germline immortality. (A) N terminal IDR sequence and schematic diagram of WAGO-1, WAGO-3, and WAGO-4 domains. IDR was predicted with metapredict^69^. (B) Animals with indicated genotype were subjected to *dpy-11* RNAi, progeny were grown on non-RNAi plates. Body length of F1 animals was quantified at L4 larval stage using ImageJ (30 animals per genotype). Student’s t test, two-tailed. (C) WT, *hrde-1(-)*, and processing-defective mutant animals expressing *gfp::h2b* in the germline (*pkIS32*) were exposed to *gfp* RNAi, after which the progeny were maintained on plates lacking RNAi. The percentage of animals expressing GFP in −1 to −3 oocytes were scored (50 animals per replicate, n = 3 biological replicates). Scale bar, 10μm. (D) Animals carrying *oma-1(zu405)* were treated with *oma-1* dsRNA and maintained at 20°C. The percentage of hatched progeny were scored across generations (n = 3 biological replicates). (E) Brood sizes of processing-defective mutant animals grown at 25°C across successive generations.

To determine whether these mutants affect TEI, we subjected them to *gfp* RNAi inheritance assays. WT animals maintained GFP silencing across generations, as expected. *wago-1(P3G)*, *wago-3(P2G, A3E)*, and *wago-4(P2G, A3E)* mutants exhibited either intact or mildly compromised inheritance, in contrast to the complete loss of TEI in *app-1(-)* and *dpf-3(-)* animals (Figures S4E-G). Similar results were observed in the *oma-1* RNAi TEI assay (Figure S4H). We next investigated whether these processing-defective mutants could recapitulate the germline immortality defects observed in *app-1(-)* and *dpf-3(-)* animals. *wago-1(P3G)* mutants became progressively sterile at 25°C (Figure S4I). Sterility was observed by the F5-F8 generation in *wago-1(P3G)* animals (Figure S4I)-2-3 generations later than the sterility seen in *app-1(-)* and *dpf-3(-)* animals (Figure 1C). *wago-3(P2G, A3E)* and *wago-4(P2G, A3E)* animals exhibited significantly reduced fertility compared with WT when reared at 25°C (Figures S4J-K). These results show that WAGO-1, WAGO-3, and WAGO-4 require proper processing by APP-1 and DPF-3 to mediate RNAi inheritance and germline immortality. These results hint that multiple substrates of APP-1 and DPF-3 must be processed correctly to promote these processes.

To test this hypothesis, we generated double and triple mutants using processing-defective point mutants of *wago-1*, *wago-3*, and *wago-4*. We then assessed whether these combined mutations phenocopied defects seen in the *app-1(-)* and *dpf-3(-)* mutants. Only double or triple mutants containing the processing-deficient *wago-1(P3G)* allele exhibited complete loss of *dpy-11* RNAi inheritance, highlighting the crucial role of WAGO-1 processing in this context (Figure 3B). In RNAi-triggered TEI assays using *gfp* and *oma-1* dsRNAs, processing-defective mutants-including *wago-1(P3G)*; *wago-3(P2G, A3E)* and the triple mutant *wago-1(P3G)*; *wago-3(P2G, A3E)*; *wago-4(P2G, A3E)*-recapitulated the TEI defects observed in the *app-1(-)* or *dpf-3(-)* mutants for both *gfp* and *oma-1* RNAi (Figure 3C-D). In contrast, double mutants such as *wago-1(P3G)*; *wago-4(P3G)* and *wago-3(P2G, A3E)*; *wago-4(P2G, A3E)* exhibited only mild TEI defects (Figure 3C-D). These results suggest that the proper processing of WAGO-1, WAGO-3, and WAGO-4 is critical for TEI. We next examined whether these mutations also affect germline immortality. Animals carrying combinations of processing-defective WAGO alleles exhibited a progressive decline in fertility, culminating in complete sterility within 4-10 generations. Notably, the triple processing mutant became sterile by generation 4, similar to *app-1(-)* and *dpf-3(-)* animals (Figure 3E). Together, these findings demonstrate that proper proteolytic processing of WAGO-1, WAGO-3, and WAGO-4 is essential for both heritable gene silencing and maintenance of germline immortality.

### Loss of predicted APP-1 and DPF-3-mediated processing leads to proteasomal degradation of WAGO-1/3/4

To investigate the effect of loss of APP-1 and DPF-3 processing on WAGO-1, WAGO-3, and WAGO-4, we performed quantitative reverse transcription PCR (qRT-PCR) to measure the mRNA levels of these three genes. The results showed that the mRNA levels of *wago-1*, *wago-3*, and *wago-4* remained largely unchanged in *app-1(-)*, *dpf-3(-)*, and *app-1(-)*; *dpf-3(-)* animals (Figure S5A). We hypothesized that APP-1- and DPF-3-mediated proteolytic processing regulates the stability of WAGO-1, WAGO-3, and WAGO-4. To test this idea, we used CRISPR/Cas9 to generate internally tagged versions of these proteins, avoiding N-terminal tagging to prevent potential processing disruptions^26^. Structural predictions indicated that WAGO-1, WAGO-3, and WAGO-4 possess intrinsically disordered, proline-rich N-terminal regions (Figure 3A)^69^. To preserve function, we inserted small HA or FLAG epitopes near the end of these regions. Functional assays confirmed that these tagged proteins largely retained activity, as evidenced by WT phenotypes in germline immortality assays (Figures S3B-E). To assess the impact of APP-1 and DPF-3 depletion on WAGO-1/3/4 protein stability, we introduced *app-1(-)*, *dpf-3(-)*, and *app-1(-)*; *dpf-3(-)* mutations into animals expressing HA- or FLAG-tagged WAGO-1/3/4. Western blot analysis revealed a significant reduction in WAGO-1, WAGO-3, and WAGO-4 protein levels upon APP-1 and/or DPF-3 loss (Figures 4A-C). Immunofluorescence and imaging revealed that the levels of WAGO-1, WAGO-3, and WAGO-4 were markedly reduced in both the shared cytoplasm and germ granules (Figures 4D-F). Processing-deficient mutants-*wago-1(P3G)::ha*, *wago-3(P2G, A3E)::ha*, and *wago-4(P2G, A3E)::ha*-also exhibited reduced protein levels compared to their WT counterparts (Figures S5B-D). These results suggest that APP-1 and DPF-3 processing is required for the stability of WAGO-1, WAGO-3, and WAGO-4.

**Figure 4.**
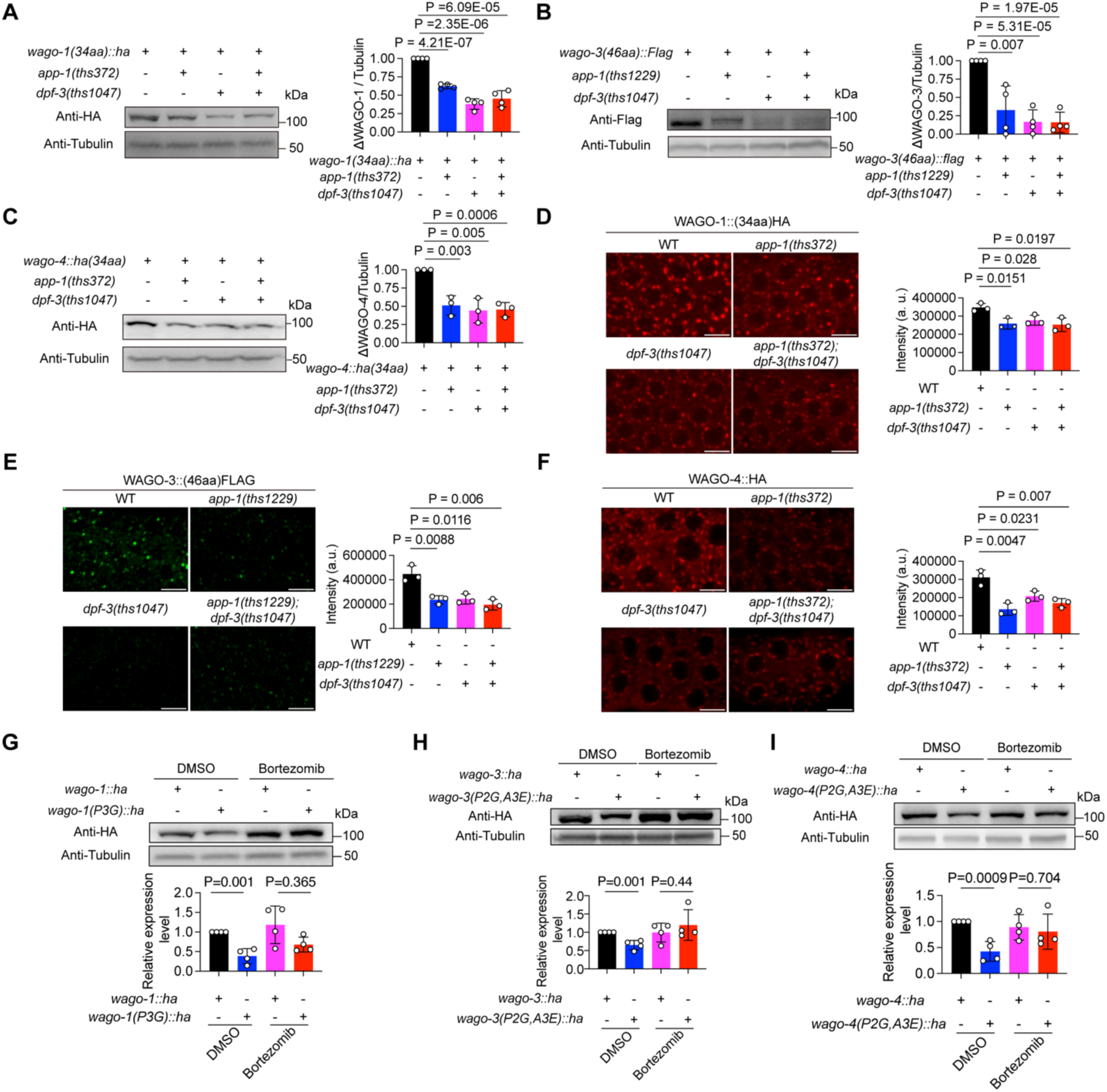
Loss of predicted APP-1- and DPF-3-mediated processing triggers proteasome-dependent degradation of WAGO-1/3/4. (A-C) Western blots showing WAGO-1, WAGO-3, and WAGO-4 protein levels in WT, *app-1(-)*, *dpf-3(-)*, and *app-1(-)*; *dpf-3(-)* animals (left). Quantification of protein abundance normalized to Tubulin is shown on the right (n = 3-4 biological replicates). Student’s t test, two-tailed. (D-F) Immunofluorescence of WAGO-1, WAGO-3, and WAGO-4 in WT, *app-1(-)*, *dpf-3(-)*, and *app-1(-)*; *dpf-3(-)* animals. Representative images of the pachytene region are shown (left), with quantification of fluorescence intensity (right, n = 3-4 animals). Student’s t test, two-tailed. (G-I) WT and processing-defective mutants were treated with DMSO or the proteasome inhibitor Bortezomib. Western blot (top) to detect WAGO-1, WAGO-3, and WAGO-4 protein level. Quantification of protein levels normalized to Tubulin (bottom, n = 4 biological replicates). Student’s t test, two-tailed.

To determine whether the reduction in WAGO-1/3/4 protein levels was due to autophagy-or proteasome-mediated degradation, we knocked down *atg-18* or *epg-5* by RNAi, both of which encode core autophagy components^70^. This treatment failed to restore WAGO protein levels (Figures S5E-J), suggesting that autophagy is not involved. However, RNAi-mediated knockdown of *spt-5*, a core proteasomal subunit, rescued WAGO-1, WAGO-3, and WAGO-4 protein levels (Figures S5K-M). Moreover, treatment with Bortezomib, a potent proteasome inhibitor^71^, restored WAGO protein levels in processing-deficient mutant animals (Figures 4G-I). Together, these findings indicate that loss of APP-1 and DPF-3 processing leads to proteasome-mediated degradation of WAGO-1, WAGO-3, and WAGO-4.

### Endogenous siRNAs reduced upon APP-1/DPF-3 loss are predominantly WAGO-1/3/4-bound siRNAs

Having established that WAGO-1/3/4 are substrates of APP-1 and DPF-3, and that APP-1/DPF-3 influence the transmission or maintenance of siRNAs during TEI, we next asked whether APP-1/DPF-3-dependent processing affects WAGO-1/3/4-associated siRNAs. We first confirmed that WAGO-1/3/4 bind siRNAs during TEI. FLAG-tagged WAGO-1, WAGO-3, and WAGO-4 were subjected to control or *oma-1* RNAi, and their associated small RNAs were sequenced. As expected, all three WAGOs co-purified with siRNAs targeting the *oma-1* locus (Figures S6A-C).

Because Argonautes stabilize their associated small RNAs^72^, we examined whether the siRNAs reduced upon APP-1/DPF-3 loss correspond to WAGO-1/3/4-bound siRNAs. Notably, the internally FLAG-tagged WAGO-1/3/4 are predicted to retain normal APP-1/DPF-3-dependent processing and function and thus should more accurately capture endogenous WAGO-1/3/4-associated siRNAs. We compared genes targeted by endogenous siRNAs associated with FLAG::WAGO-1, FLAG::WAGO-3, and FLAG::WAGO-4 with those targeted by siRNAs decreased in *app-1(−)*, *dpf-3(−)*, and *app-1(−)*; *dpf-3(−)* mutants (Figure S6D). Strikingly, siRNAs reduced upon APP-1/DPF-3 loss were significantly enriched for WAGO-1/3/4-bound species: 84%, 84%, and 80% of decreased siRNAs in the three mutants, respectively, corresponded to WAGO-1/3/4-associated siRNAs (Figure S6E). The reduction in these siRNAs may result from proteasome-mediated degradation of WAGO-1/3/4-siRNAs complex or increased siRNA degradation due to the reduced WAGO-1/3/4 pool. Together, these findings indicate that loss of APP-1/DPF-3 processing selectively decreases WAGO-1/3/4-bound siRNAs, thereby contributing to defects in TEI and germline immortality.

### The conserved E3 Ligases RNF-1 and UBR-7 mediate the degradation of unprocessed WAGO-1, WAGO-3, and WAGO-4

E3 ubiquitin ligases recognize proteasome substrates and catalyze the attachment of ubiquitin chains, targeting bound proteins for proteasomal degradation^73^. To identify E3 ligases that regulate the stability of unprocessed WAGO-1, WAGO-3, and WAGO-4, we depleted *rpt-5* with RNAi and immunoprecipitated FLAG-tagged WAGO-1, WAGO-3, and WAGO-4 in *app-1(-)*; *dpf-3(-)* or *app-1(-)* background using anti-FLAG magnetic beads. Co-precipitating proteins were analyzed by mass spectrometry. As expected, WAGO-1, WAGO-3, and WAGO-4 were among the most enriched proteins in their respective immunoprecipitants (Table S2-5). Notably, we also detected several proteasome subunits (For instance, RPT-1, RPT-3, and RPT-6) and six conserved, predicted E3 ubiquitin ligases-RNF-1, C38D9.2, UBR-7(T22C1.1), RFP-1, RBPL-1, and SUT-2-among the associated proteins (Table S2-5). To identify which of these E3 ligases might directly interact with WAGO-1/3/4, we performed yeast two-hybrid assays (Figure 5A). These assays revealed that only RNF-1 ligases interacted with all three WAGOs and UBR-7 may interact weakly with WAGO-3 and WAGO-4 (Figures 5B-C, S7A-I). To test whether these E3 ligases regulate WAGO-1/3/4 stability, we depleted each E3 ligase by RNAi in *app-1(-)*; *dpf-3(-)* animals and examined WAGO-1/3/4 protein levels by immunofluorescence. Surprisingly, knockdown of *ubr-7* largely restored WAGO-1, WAGO-3, and WAGO-4 protein levels, whereas knockdown of *rnf-1* largely rescued WAGO-3 and WAGO-4 but not WAGO-1(Figures 5D-F, S8A-F). Because RNAi may not fully eliminate RNF-1, we generated a *rnf-1(-)* mutant, in which depletion of RNF-1 largely restored all three WAGOs (Figures 5G-L). Given that both RNF-1 and UBR-7 influence WAGO-1/3/4 stability, we next asked whether these E3 ligases interact with each other. Indeed, yeast two-hybrid analysis confirmed a direct interaction between RNF-1 and UBR-7 (Figure 5M). To determine the subcellular localization of RNF-1 and UBR-7, we used CRISPR/Cas9 to insert GFP tags at their endogenous loci. Fluorescence imaging revealed that GFP::RNF-1 was distributed throughout the cytoplasm of the *C. elegans* germline and embryos (Figure S8G). Unexpectedly, UBR-7::GFP localized predominantly to the nucleus in the germline. In embryos, UBR-7::GFP fluorescence was detected in both the cytoplasm and the nucleus (Figure S8H). Because E3 ligases can be co-degraded with their substrates by the proteasome, we next asked whether inhibiting proteasome activity by knocking down *rpt-5* would increase RNF-1 and UBR-7 abundance. Indeed, GFP::RNF-1 and UBR-7::GFP levels were elevated following rpt-5 RNAi treatment. Notably, UBR-7::GFP fluorescence became apparent in the cytoplasm after *rpt-5* RNAi (Figures S8G and S8H). Together, these results support a model in which RNF-1 recognizes WAGO-1/3/4 and cooperates with UBR-7 to mediate their ubiquitination and degradation in the cytoplasm of the *C. elegans* germline.

**Figure 5.**
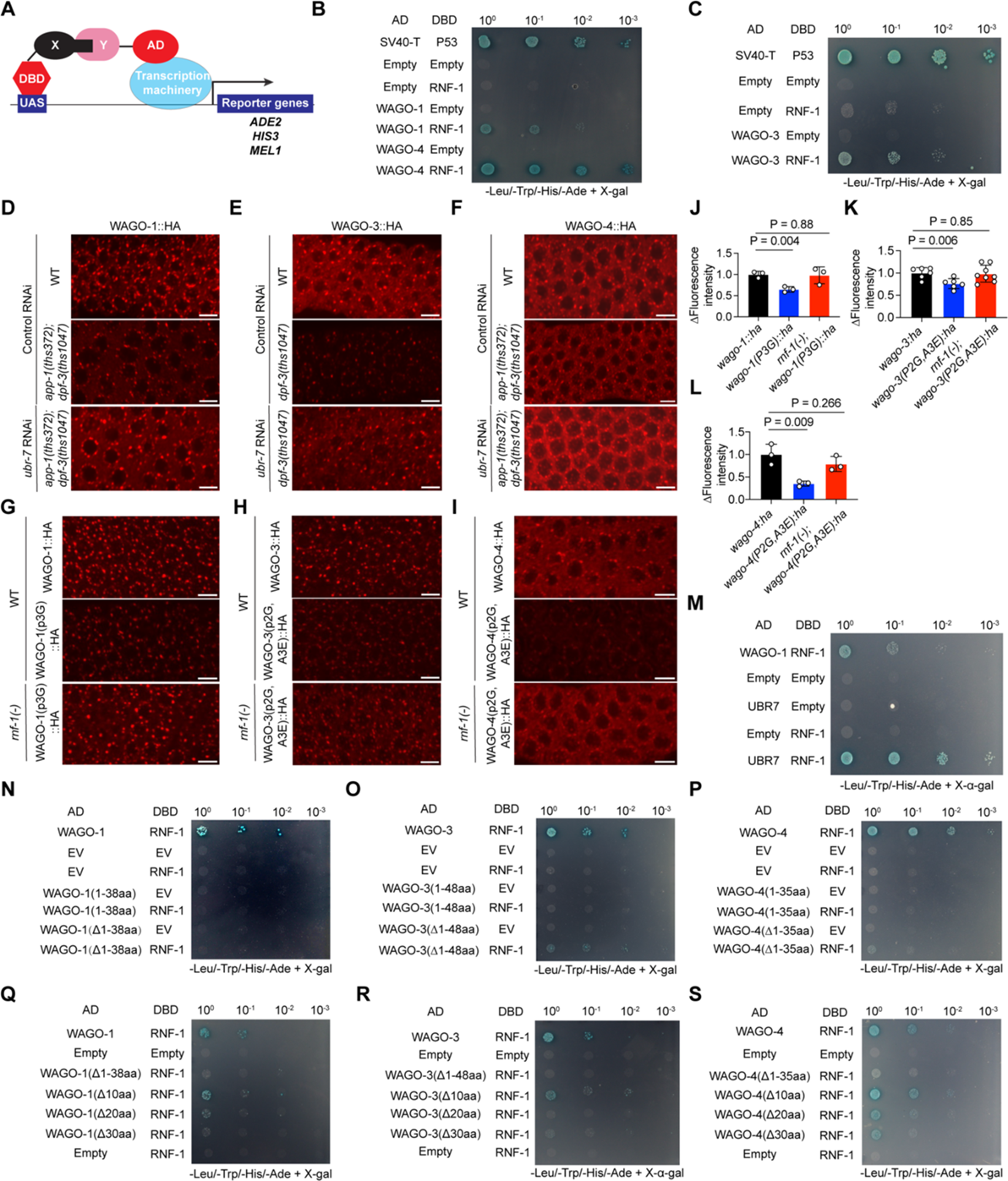
Identification of E3 ligases that mediate degradation of WAGO-1, WAGO-3, and WAGO-4. (A) Schematic of the yeast two-hybrid system used in this study. Yeast cells co-transformed with plasmids expressing GLA4 AD- and DBD-fused proteins grow on plates lacking Leu and Trp. Interaction between proteins X and Y activates the reporter genes *ADE2*, *HIS3*, and *MEL1*, enabling growth on plates lacking Ade and His and producing blue colonies in the presence of X-gal. (B-C) Yeast two-hybrid assays testing the interaction between RNF-1 and WAGO-1/3/4. The interaction between SV40 T antigen and p53 was used as a positive control^95^. (D-F) Immunofluorescence imaging of WAGO-1/3/4 in WT and *app-1(-)*; *dpf-3(-)* animals treated with control RNAi or *ubr-7* RNAi. Scale bar, 5μm. (G-I) Immunofluorescence imaging of WT WAGO-1/3/4 and their N-terminal processing mutants in WT and *rnf-1(-)* background. Scale bar, 5μm. (J-L) Quantification of the relative fluorescence intensity of WAGO-1/3/4 shown in (G-I). n = 3-6 animals. Student’s t test, two-tailed. (M) Yeast two-hybrid assay testing the interaction between RNF-1 and UBR-7. n = 3 biological replicates. (N-P) Yeast two-hybrid assay testing the interaction between RNF-1 and the WAGO-1/3/4 IDR or WAGO-1 lacking its IDR. n = 3 biological replicates. (Q-S) Yeast two-hybrid assays testing the interaction between RNF-1 and WAGO-1/3/4 variants lacking 10 aa, 20 aa, or 30 aa of their IDRs. n = 3 biological replicates.

To investigate how RNF-1 interacts with unprocessed WAGO-1/3/4, we performed yeast two-hybrid assays using a series of WAGO-1/3/4 truncations. First, no interactions were detected between RNF-1 and IDRs of WAGO-1/3/4, suggesting that RNF-1 does not directly recognize the unprocessed N-terminal residues (Figures 5N-P, S7M-O). Second, deletion of the IDRs of WAGO-1/3/4 abolished the interaction between RNF-1 and each of the three WAGOs (Figures 5N-P, S7M-O). Third, we removed 10aas, 20aas, and 30aas from WAGO-1/3/4 IDRs and performed similar yeast two hybrid assays. The results showed that loss of N terminal 10aa-20aa decreased the interactions between WAGO-1/3/4 and RNF-1 (Figures 5Q-S, S7P-R). Together, these results show that the interaction between RNF-1 and WAGO-1, WAGO-3, and WAGO-4 is regulated by length of their N-terminal IDRs, and these results hint that RNF-1 may recognize conformational changes in WAGO-1/3/4 controlled by their N-terminal IDRs (see Discussion).

### N terminal IDR of WAGO-1, WAGO-3, and WAGO-4 were processed *in vivo* and are blocked by loss of APP-1 and DPF-3

To determine whether APP-1 and DPF-3 process WAGO-1/3/4 in vivo, we used mass spectrometry to map the N termini of endogenous, internally FLAG-tagged WAGO-1, WAGO-3, and WAGO-4 proteins immunoprecipitated from *app-1(−)*, *dpf-3(−)*, and *app-1(−)*; *dpf-3(−)* mutant animals. Multiple N-terminal species of WAGO-1 and WAGO-3 with different lengths were detected (Figures 6A and 6B). For WAGO-4, a processed species lacking the first six amino acids was detected (Figure 6C). Processing of WAGO-1 and WAGO-4 was abolished in *app-1(−)*, *dpf-3(−)*, and *app-1(−)*; *dpf-3(−)* mutant animals (Figures 6A and 6C).

**Figure 6.**
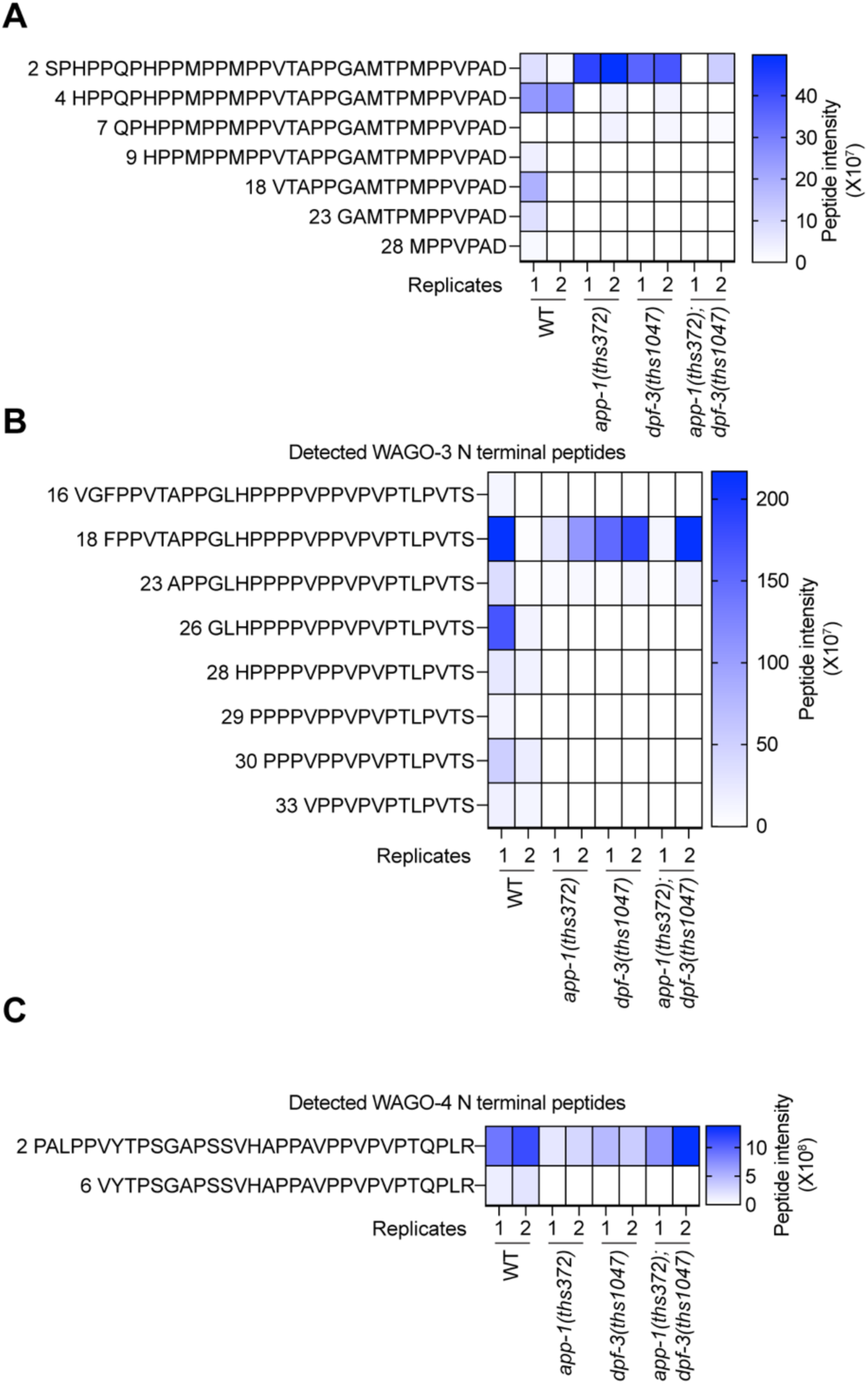
Processing of WAGO-1/3/4 in vivo is impaired by loss of APP-1 and DPF-3. (A-C) endogenous FLAG-WAGO-1/3/4 were immunoprecipitated, N terminal processed species were detected by Mass Spectrometry.

For WAGO-3, we failed to detect the full-length N terminus, possibly due to N-terminal degradation or because the predicted N-terminal fragment (46–50 amino acids) was below the detection limit of mass spectrometry (Figure 6B). Instead, we detected a species lacking the first 18 amino acids, for reasons that remain unclear (Figure 6B). Notably, other N-terminally processed species of WAGO-3 were not detected in *app-1(−)*, *dpf-3(−)*, or *app-1(−)*; *dpf-3(−)* mutant animals (Figure 6B). Together, these results indicate that loss of APP-1 and DPF-3 largely suppresses the processing of WAGO-1, WAGO-3, and WAGO-4.

## Discussion

Small RNA-based gene regulation is a deeply conserved mechanism, at the core of this is the Argonautes that associate with small RNAs. All Argonautes share conserved PAZ, MID domain that anchor the 3’ and 5’ ends of small RNAs, respectively and PIWI domain that is structually similar to RNaseH, while their N-terminal IDRs are highly divergent and thought to play regulatory roles^74,75^. Previous studies have shown that the Argonaute N-terminus influences protein stability and small RNA duplex unwinding^76,77^. Here, we identify cooperative N-terminal processing by two peptidases that trim the IDRs of three germline-expressed Argonautes-WAGO-1, WAGO-3, and WAGO-4. This processing appears to have two major effects: it enhances WAGO protein stability and proper siRNA loading (Ida Isolehto *et al*, parallel submission) in the germ granules subcompartments (Figure 7). Notably, unprocessed WAGO-1, WAGO-3, and WAGO-4 interact with the conserved E3 ligase RNF-1, which recruits another conserved E3 ligase, UBR7, to promote their proteasome-mediated degradation. However, RNF-1 does not bind the isolated IDRs of WAGOs. We speculate that IDRs modulate conformational states of WAGO-1/3/4 that are recognized by RNF-1. In this presumptive conformational state, the IDR occupies the siRNA-binding groove when siRNAs are not loaded (Ida Isolehto *et al*, parallel submission).

**Figure 7.**
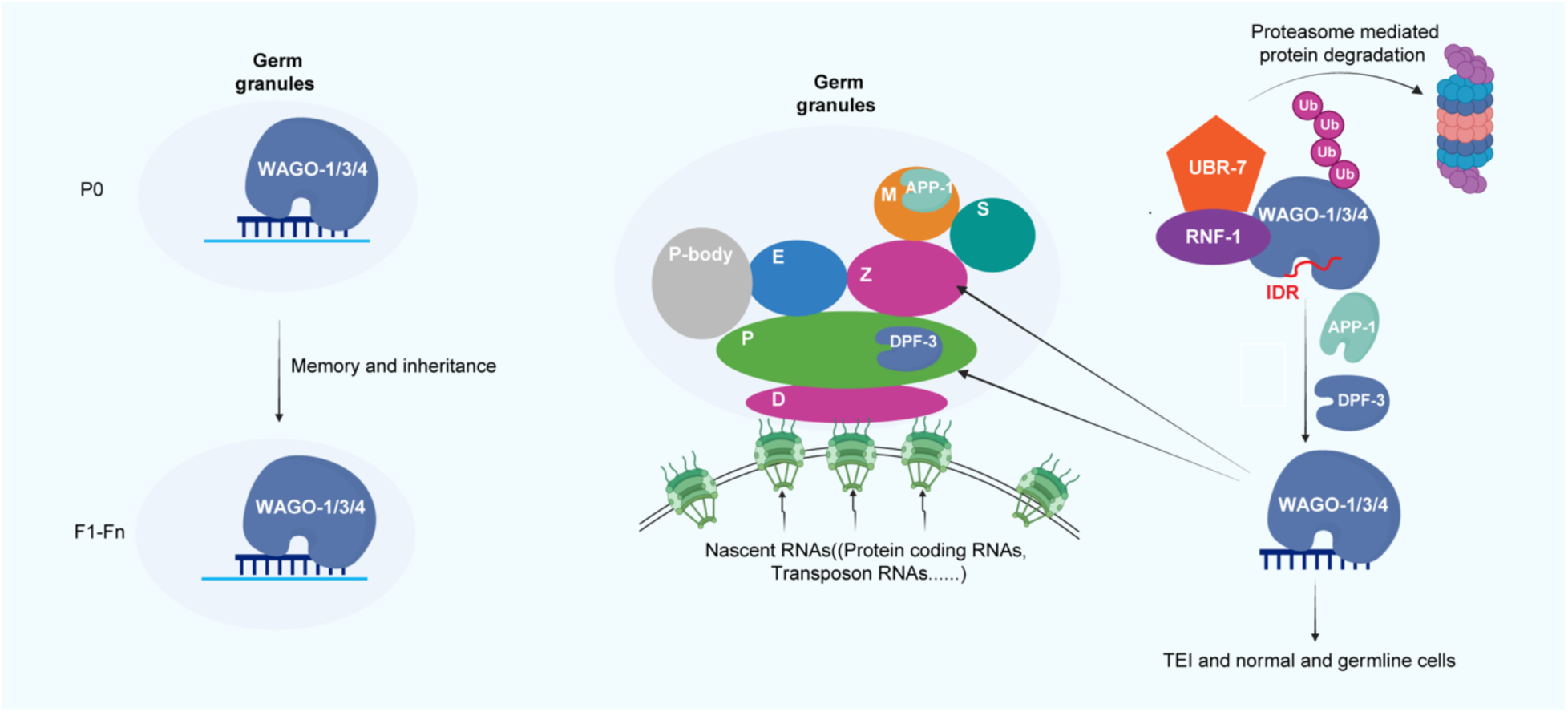
Working model. WAGO-1/3/4–bound siRNAs help maintain epigenetic memory within perinuclear germ granules. The N-terminal IDRs of WAGO-1/3/4 can occupy their own siRNA-binding grooves, adopting a conformation recognized by the E3 ligase RNF-1, which in turn recruits the E3 ligase UBR-7 to ubiquitinate WAGO-1/3/4 and target them for proteasome-mediated degradation. APP-1 and DPF-3 cooperatively process the N-terminal IDRs of WAGO-1/3/4, thereby preventing RNF-1–dependent recognition and stabilizing WAGO-1/3/4 within germ-granule subcompartments. This processing promotes the accumulation of WAGO-1/3/4, enabling efficient TEI and supporting normal germline immortality.

Although APP-1 and DPF-3 are enriched in distinct germ granule subcompartments-the Mutator foci and P granules, respectively-a fraction of both enzymes localizes to the shared rachis region of the germline. This raises the possibility that the cooperative processing of WAGO-1, WAGO-3, and WAGO-4 occurs in this shared cytoplasmic space before Argonautes are partitioned into specific germ granule subdomains. Alternatively, cooperative processing might still take place within granule subcompartments during transient interactions between the Mutator foci and P granules at defined developmental stages^78^. In vivo processing reveals that WAGO-1 and WAGO-3 are trimmed by approximately 20–30 amino acids within their N-terminal IDRs. These observations raise the possibility that the high local concentration of APP-1 and DPF-3 within germ granules enhances their processivity and catalytic activity. Perhaps the low activity in the diffused cytoplasm prevents WAGOs from processing even though APP-1 and DPF-3 are present, thus preventing abbrent loading siRNAs for essential genes such as histone genes (Ida Isolehto *et al*, parallel submission), while efficient processing in the germ granules promote the stability thus accumulation of WAGOs as well as siRNA loading in the germ granules. This added another layer of spatiotemporal regulation of small RNA-based gene regulation. The processing may prevent unprocessed WAGOs from aberrantly engaging small RNA machineries and interfering with gene silencing. The E3 ligase-proteasome degradation system uncovered here likely acts as a quality control mechanism that eliminates improperly processed Argonautes.

The substrate specificities of APP-1 and DPF-3 may extend beyond their canonical motifs^79,80^. DPF-3 has been shown to cleave after residues such as XaaT, XaaS, and XaaG and may even have tripeptidly peptiese activity^79–81^, while the proteome-wide specificity of APP-1 remains undefined but may reach beyond XaaP. Although cleavage of noncanonical substrates is likely less efficient-potentially constrained by steric factors-the processing efficience maybe enhanced by enrichment of APP-1 and DPF-3 in the corresponding germ granule subcompartments.

APP-1 and DPF-3 are highly conserved across species, with mammalian homologs XPNPEP1 and DPP8/9, respectively^52,82^. This conservation suggests that the cooperative processing we describe may represent a broadly conserved mechanism regulating diverse substrates and biological processes. In silico proteome-wide predictions in *C. elegans* and humans identified 176 and 328 proteins, respectively, that may be cooperatively processed by APP-1/DPF-3 or XPNPEP1 and DPP8/9 based on canonical processing motifs. Gene enrichment analysis highlighted processes such as transcriptional regulation and immunity among the most affected categories (Figures S9A-B). Indeed, XPNPEP1 and DPP8/9 have been implicated in the cooperative regulation of inflammasome activation^83,84^. Notably, the human Xaa-Pro aminopeptidase family has three members, and the dipeptidyl peptidase family has ten, yet most of their substrate specificities remain poorly defined at the proteome scale^48,54^. N-terminal residues play critical roles in determining protein localization, stability, and modification status through mechanisms such as resistance to peptidase hydrolysis, the N-degron pathway, and post-translational modifications including methylation and acetylation^50^. We propose that cooperative N-terminal processing may intersect with these pathways, influencing ubiquitin–proteasome and autophagy-mediated degradation and ultimately determining protein fate. Intriguingly, while N-terminal proteolysis is generally thought to reduce protein stability, our findings reveal the opposite effect-N-terminal trimming can stabilize Argonautes and promote their functional maturation. Future systematic mapping of N-terminal diversity across the proteome will be crucial to determine how widespread such cooperative protease processing is, and how it shapes protein stability and regulatory networks in diverse biological contexts.

## Materials and Methods

### Strains

All *C. elegans* strains(Table S1) except *oma-1(zu405)* containing strains (maintained at 15°C) were maintained at 20°C and cultured on NGM plates seeded with E. coli OP50, following previously described methods^85^, unless otherwise specified. Certain strains were obtained from the CGC (P40 OD010440). “WT” refers to the N2 Bristol strain unless stated otherwise. All strains are available upon request.

### CRISPR/Cas9 mediated gene editing

We employed the co-CRISPR strategy to generate deletions or insertions at the endogenous locus^25,86,87^. The sgRNAs were synthesized via *in vitro* transcription using T7 RNA polymerase. Repair templates containing ~100bp homology arms (for HA/FLAG tagging or point mutations) or ~400bp homology arms (for GFP insertion) were generated by PCR amplification and reannealed using the following thermocycling program: 95°C for 2 minutes, 85°C for 10 seconds, 75°C for 10 seconds, 65°C for 10 seconds, 55°C for 1 minute, 45°C for 30 seconds, 35°C for 10 seconds, and 25°C for 10 seconds. A mixture of 1.8μg purified sgRNAs targeting the genes of interest and *unc-58* was incubated with 5μg purified 3×NLS-SpCas9 at 37°C for 15 minutes^25^. Repair templates, including 24ng/μL AF-JA-76 and 25ng/μL *gfp*, *ha*, *flag*, or point-mutation repair templates, were then added to generate a 10μL injection mixture. This mixture was injected into the gonads of young adult animals. The injected animals were maintained at 20°C for three days, after which uncoordinated (unc) progeny were selected and genotyped. Deletions were generated using two sgRNAs located near the start and stop codons without the use of repair templates. Domain and allele schematics were generated using IBS 2.0^88^. Detailed information on the sgRNAs used in this study can be found in Table S2.

### RNAi and RNAi inheritance

A single clone of freshly streaked Escherichia coli HT115 bacteria containing plasmids expressing either control dsRNA (L4440) or dsRNAs targeting specific genes (Table S3) was grown in Luria-Bertani (LB) broth overnight for 16 hours. Expression of dsRNAs was induced by adding 4mM IPTG at room temperature for 2 hours before seeding the cultures onto RNAi plates (NGM plates containing 1mM IPTG and 25µg/mL carbenicillin). Gravid adults were treated with hypochlorite to isolate embryos (egg preparation).The isolated embryos were then transferred to the RNAi plates. To study RNAi inheritance, egg preparation was performed on P0 animals previously exposed to RNAi treatment, and the resulting embryos were placed on either OP50 plates (no RNAi) or L4440 plates. For *gfp* RNAi inheritance, GFP expression was scored under a 10× objective, and the percentage of GFP-expressing animals was scored. For *dpy-11* RNAi inheritance, the body length of L4 larval-stage animals in the inheriting F1 generation was measured using ImageJ. For *oma-1* RNAi inheritance, six young adult animals were randomly selected to lay 100–200 embryos, after which they were removed. The percentage of hatched embryos was then assessed the following day. For RNAi targeting *rpt-5*, early L4 larval stage animals were transferred to RNAi plates containing HT115 bacteria expressing *rpt-5* dsRNAs and allowed to grow for 36 hours before being harvested for downstream analysis. For RNAi targeting candidate E3 ligases, the isolated embryos were grown on RNAi plates for 3 days before immunofluorescence at young adult stage.

### Mrt assay

Six L4 larval-stage animals raised at 20°C were randomly selected and transferred to 25°C. In subsequent generations, six L4 larvae were randomly picked and maintained at 25°C until the population became sterile. The progeny of these six animals were scored at the specified generations. To determine whether the germline immortality defect is due to epigenetic changes, six L4 larvae were randomly selected and raised at 25°C until the population was nearly sterile. Six L4 larvae progeny were then transferred to 20°C and allowed to propagate for specified generations until their brood size was restored. This process was repeated by transferring six L4 larvae alternately between 25°C and 20°C. The progeny of these six animals were scored at the specified generations.

### Bortezomib treatment

Bortezomib treatments were carried out as previously described^71^. Young adult animals (1000–2000) were collected and suspended in 1ml S basal buffer (1g/L K_2_HPO_4_, 6g/L KH_2_PO_4_, 5mg/L cholesterol, and 5.85g/L NaCl), supplemented with OP50 (equivalent to 4 mL OP50), 50μg/mL Carbenicillin, and either 0.01% DMSO (control) or the desired concentration of bortezomib (5μg/mL or 10μg/mL), which was diluted from a 50mg/mL stock in DMSO. The suspension was placed into 6-well tissue culture plates, with a final volume of 1ml per well. The plates were incubated at 20°C for 8 hours before being harvested for Western blot analysis.

### Immunofluorescence

Young adult *C. elegans* were placed on a SuperFrost Plus slide (1255015, Thermo Fisher Scientific) and dissected in 1× egg buffer (25mM HEPES pH 7.3, 118mM NaCl, 48mM KCl, 2mM CaCl_2_, and 2mM MgCl_2_) using a 1mL syringe needle. After dissection, a coverslip was gently placed on top, and excess buffer was removed using a Kimwipe. The slide was then flash-frozen by placing it on a pre-cooled metal block at −80°C for 10 minutes. Once frozen, the coverslip was carefully removed with a razor blade, and the slide was immediately transferred to pre-cooled methanol (−20°C) for 10 minutes. Following methanol fixation, the slide was washed twice with 1× PBST (1× phosphate-buffered saline with 0.1% Tween-20). The samples were then fixed with 4% paraformaldehyde in PBS. A piece of parafilm (cut to the size of the coverslip) was placed over the samples, and the slide was incubated at room temperature for 20 minutes. After fixation, the slide was washed twice with 1× PBST and incubated overnight at room temperature in a humid chamber with 100μL of diluted primary antibody. The next day, the slide was washed three times with 1× PBST, followed by incubation with 100μL of diluted secondary antibody for 2 hours at room temperature in a humid chamber. Afterward, the slide was washed three more times with 1× PBST. Finally, the samples were mounted using VECTASHIELD Antifade Mounting Medium containing 1μg/mL DAPI (H-1000-10; Vector Labs). A coverslip was carefully placed on top and sealed with nail polish to preserve the samples. The following primary antibodies were used: anti-HA antibody (1:250; Cell Signaling Technology, 3724S), anti-FLAG antibody (1:500; Sigma-Aldrich, F1804) and anti-PGL-1 antibody (1:20; Developmental Studies Hybridoma Bank, cloneK76, RRID: AB_531836). The following secondary antibodies were used: goat anti-mouse IgG (H+L) cross-adsorbed secondary antibody, Alexa Fluor™ 488 (1:50; Thermo Fisher Scientific, A11001) and goat anti-rabbit IgG (H+L) cross-adsorbed secondary antibody, Alexa Fluor™ 568 (1:50; Thermo Fisher Scientific, A11011).

### Microscopy

L4 larval animals or Day 1 young adult *C. elegans* were anesthetized with either 0.1% sodium azide or 0.25μM levamisole hydrochloride in M9 buffer and mounted on a glass slide. The animals were then imaged immediately using a NIKON ECLIPSE Ti2-E microscope equipped with a CSU-570 W1 spinning disc and a Plan Apochromat 100×/1.45 oil immersion objective. For animals expressing GFP::H2B, the same preparation procedure was followed, and imaging was conducted promptly using a Leica DMI8 inverted microscope with a 63×/1.4 Plan Apo oil immersion objective.

### Colocalization analysis with ImageJ

To evaluate the degree of colocalization between two granules, an auto-threshold value was determined for each granule. This threshold value was applied to the three-dimensional (3D) segmentation function in the ImageJ 3D Manager plug-in, which incorporated the granule images into the 3D Manager. A white mask was then generated around the two granules using the Coloc2 plug-in. The degree of colocalization, quantified by Pearson’s R value, was calculated using Coloc2 in Fiji.

### qRT-PCR

Total RNA was extracted with RNAiso Plus (Takara, 9109) and complementary DNA (cDNA) was synthesized using the HiScript III 1st Strand cDNA Synthesis Kit (+gDNA wiper, Vazyme, R312-02) according to vendor’s instructions. mRNA levels were quantified using the SYBR green Pro Taq HS kit (Accurate Biology, AG11701). The qRT-PCR reactions were run on a Roche LightCycler 480 II. Primer sequences are listed in Table S4.

### FLAG immunoprecipitation and mass spectrometry (FLAG IP-Mass spec)

L4-stage larvae were treated with *rpt-5* RNAi for 36h. Approximately 80,000 young adult worms were collected for each strain: *app-1(ths372)*; *dpf-3(ths1047)*, *app-1(ths372)*; *dpf-3(ths1047)*; *flag::wago-4*, *app-1(ths372)*; *dpf-3(ths1047)*; *flag::wago-3*, and *app-1(ths372)*; *flag::wago-1*. Worms were flash-frozen by dripping dropwise into liquid nitrogen and stored at −80°C. Frozen animals were ground to a fine powder under liquid nitrogen and resuspended in 1mL of 1× lysis buffer (50mM Tris-HCl, pH 7.5, 150mM NaCl, 0.25% Triton X-100, 1 mM EDTA) supplemented with 1 mM PMSF and 1× Complete protease inhibitor cocktail (Roche; 11 873 580 001). The suspension was rotated for 1h at 4°C. Lysates were clarified by centrifugation at 17,949 × g for 15min at 4°C, and the supernatants were filtered through a 0.45μm filter unit (Millipore; SLHP033RS). Protein concentrations were determined using the BCA assay. For each sample, 5 mg of total protein was incubated with 50μL of 50% slurry anti-FLAG M2 magnetic beads (Sigma-Aldrich; M8823) for 2h at 4°C. Beads were washed once with lysis buffer containing protease inhibitors, followed by six washes with TBS buffer (50mM Tris-HCl, pH 7.5, 150mM NaCl) without protease inhibitors. Bound proteins were eluted by rotating the beads in 500μL of 500mM NH_4_OH for 20min at 37°C.

Samples were subjected to tryptic digestion using a commercial reagent kit (B0770-20C; KE FU TECHNOLOGY) according to the manufacturer’s instructions. Liquid chromatography-tandem mass spectrometry (LC–MS/MS) analysis was performed on an UltiMate 3000 RSLC nano system coupled to an Orbitrap Fusion Lumos Tribrid mass spectrometer (Thermo Fisher Scientific). Peptides were loaded onto a C18 pre-column (Acclaim PepMap; 75 μm × 2 cm; nanoViper; 3 μm; 100 Å) and separated on a C18 analytical column (Acclaim PepMap; 75 μm × 15 cm; nanoViper; 2 μm; 100 Å; Thermo Fisher Scientific). Peptide separation was achieved using a linear gradient of 3-45% buffer B (0.1% formic acid and 20% water in acetonitrile) over 55min, followed by 45–98% buffer B for 5min and a hold at 98% buffer B for 5min, at a constant flow rate of 300nL min⁻¹ (buffer A: 0.1% formic acid in ultrapure water). Survey scans were acquired in positive ion mode over an m/z range of 350–1,500 at a resolution of 60,000 (full width at half maximum at m/z 200). Data-dependent acquisition was performed with a cycle time of 3s. Precursor ions were fragmented by higher-energy collisional dissociation with 30% normalized collision energy, and fragment ions were detected at a resolution of 15,000. Dynamic exclusion was enabled for 60s with a 10-ppm mass tolerance around the precursor and its isotopes. Raw mass spectrometry files (Thermo.raw format) were processed using MaxQuant (v1.6.2.10)^89^ for peptide identification against the *C. elegans* UniProtKB/Swiss-Prot database (downloaded on 6 February 2024). Each experiment was performed with two biological replicates, and the “match between runs” option was enabled for data integration. Trypsin was specified as the digestion enzyme, allowing up to two missed cleavages. Carbamidomethylation of cysteine was set as a fixed modification, while N-terminal acetylation and methionine oxidation were included as variable modifications. Label-free quantification (LFQ) was performed using the fast LFQ algorithm with a minimum ratio count of 3 and an average of 6, based on the iBAQ approach. ProteinGroups.txt files generated by MaxQuant were filtered to retain proteins identified by at least two unique peptides in both replicates. The fraction of total (FOT) for each protein was calculated as: FOTn = iBAQn/∑iBAQ, where n represents each protein.

### Western blot

Young adult animals were allowed to settle, after which the supernatant was removed. An equal volume of worm boiling buffer (100mM Tris-HCl, pH 6.8, 4% SDS, 20% glycerol) was added. Samples were incubated at 100°C for 8min, vortexed, and reheated for an additional 8min. Lysates were clarified by centrifugation at 10,000 × g for 5min, and the supernatants were resolved by SDS–PAGE and transferred to PVDF membranes. Membranes were probed overnight at 4°C with the following primary antibodies: anti-HA (1:1000; CST, 3724S), anti-FLAG M2 (1:1000; Sigma-Aldrich, F1804), anti-tubulin (1:5000; RayBiotech, RM2003), and anti-FLAG M2–peroxidase (HRP) (1:1000; Sigma-Aldrich, A8592). The following secondary antibodies were used: goat anti-rabbit HRP (1:2000; RayBiotech, RM3002) and goat anti-mouse HRP (1:2000; RayBiotech, RM3001).

### Small RNA RIP-seq and small RNA sequencing from total RNAs

RNA immunoprecipitations were done as described previously^19,90^. Briefly, 20,000 young adults were collected and flash frozen in liquid nitrogen. Samples were resuspended in sonication buffer (20mM Tris-HCl pH 7.5, 200mM NaCl, 2.5mM MgCl2,10% glycerol, 0.5% NP-40, 160U/ml RiboLock RNase Inhibitor, 1mM DTT and Roche protease inhibitor cocktail without EDTA) and sonicated for 6 cycles (30s on, 30s off at low power at 4°C) using a Biorupter sonicator (Diagenode). Lysates were clarified by centrifugation at 20817g for 15min. Supernatants were incubated with Anti-FLAG M2 magnetic beads (Sigma-Aldirch, M8823) for 2h at 4°C. Beads were washed six times with RNA immunoprecipitation buffer (20mM Tris-HCl pH 7.5, 200mM NaCl, 2.5mM MgCl2, 10% glycerol, 0.5% NP-40). Protein and associated RNAs were eluted with 100mg/ml 3xFLAG peptide (MedChemExpress, HY-P0319). RNAs were extracted with TRIzol reagent followed by precipitation with 2.5X volume ethanol. Purified RNAs were treated with RNA 5’ Polyphosphatase (Lucigen, RP8092H) to remove 5’-di-phosphorylation. Small RNAs were then cloned with VAHTS Small RNA Library Prep Kits (Vazyme, NR811) kits following vendor’s instructions. Cloned libraries were sent to HaploX company for high through sequencing.

For cloning small RNAs from total RNAs, total RNAs from WT, *app-1(-)*, *dpf-3(-)*, *app-1(-); dpf-3(-)* young adult animals were extracted with TRIzol reagent. Subsequently, the total RNAs were treated with RNA 5’ Polyphosphatase (Lucigen, RP8092H) to remove 5’-di-phosphorylation. Small RNAs were then cloned with VAHTS Small RNA Library Prep Kits (Vazyme, NR811) following vendor’s instructions. Cloned libraries were sent to HaploX company for high through sequencing.

### Bioinformatic analysis of small RNA-seq data

Raw sequencing data were first assessed using FastQC. R1 reads were extracted and subjected to adapter trimming with Cutadapt (parameters: -m 16 -M 30 --discard-untrimmed) to remove the adapter sequence AGATCGGAAGAGCACACGTCTGAACTCCAGTCA. Trimmed reads were re-evaluated with FastQC to confirm the quality of the remaining insert sequences. After quality control, reads were aligned to the *C. elegans* reference genome WS292 using Bowtie^91^ (parameters: -f -v 0 -k 5 --best --strata). Alignment files were converted to BAM format using Samtools. Gene-level read counts were generated with featureCounts (-s 2)^92^. Annotation for repetitive elements were obtained using the RepeatMasker (rmsk) corresponding to the *C. elegans* ce11 assembly (downloaded from the UCSC). Differential expression analysis was performed using the DESeq2^93^ package in R. Log2 fold changes were estimated by the DESeq2 model, and P values were adjusted for multiple testing using the Benjamini-Hochberg procedure to obtain adjusted P values. Representative factors were calculated using the online tool “statistical significance of the overlap between two groups of genes” (http://www.nemates.org/MA/progs/overlap_stats.html), assuming a total of 46,927 genes in the WS292 genome. All scripts and software version information used in the analysis are available upon request.

### Yeast two hybrid

Full-length or truncated cDNAs of *wago-1*, *wago-3*, *wago-4*, and candidate E3 ligases were cloned into the EcoRI and BamHI sites of either pGADT7 or pGBKT7 vectors. The yeast strain Y2HGold was transformed with 1µg of plasmid DNA using carrier DNA, lithium acetate, and PEG3350, following previous reported protocols^94^. Transformants were selected on SD/-Leu/-Trp plates at 30°C to confirm the presence of both plasmids. Protein-protein interactions were assessed by plating on SD/-Leu/-Trp/-Ade/-His selective media supplemented with 0.04mg/ml X-α-Gal (prepared by adding 1ml of 20mg/ml X-α-Gal to 500ml of SD/-Leu/-Trp/-Ade/-His medium). Plates were incubated at 30°C for three days, and yeast growth and color development were documented photographically.

### *In vivo* protease processing and mass spectrometry

Approximately 20,000 young adult worms were collected and flash-frozen in liquid nitrogen. Samples were resuspended in 1mL of lysis buffer (50mM Tris-HCl, pH 7.5, 150mM NaCl, 0.25% Triton X-100, 1mM EDTA) supplemented with 1 mM PMSF and 1× Complete protease inhibitor cocktail (Roche, 11 873 580 001), and lysed by sonication using a Bioruptor sonicator (Diagenode) for 8 cycles (30s on, 30s off) at high power at 4°C. Lysates were clarified by centrifugation at 20,817g for 1min at 4°C and filtered through a 0.45μm filter unit (Millipore, SLHP033RS). The protein concentrations were determined by BCA assay. For immunoprecipitation, 5mg of total protein was incubated with anti-FLAG M2 magnetic beads (Sigma-Aldrich, M8823) for 2h at 4°C with rotation. Beads were washed once with lysis buffer containing protease inhibitors, followed by six washes with TBS buffer (50mM Tris-HCl, pH7.5, 150mM NaCl) without protease inhibitors. On-bead digestion was performed essentially as described previously^48,81^. Briefly, beads were resuspended in 5μL of digestion buffer (3M guanidine hydrochloride, 20mM EPPS, pH 8.5, 10mM 2-chloroacetamide (CAA), 5mM TCEP), followed by addition of 1μL Lys-C (0.2μg/μL in 50mM HEPES, pH 8.5). Samples were incubated at 21°C for 4 h with shaking, after which 17μL of 50 mM HEPES (pH 8.5) was added. Trypsin (1μL at 0.2μg/μL) was then added and samples were incubated overnight at 37°C, followed by addition of a further 1μL of trypsin and incubation for an additional 4h. For WAGO-3 samples, Asp-N protease (1μL at 0.2 μg/μL) was added simultaneously with trypsin to enable generation of an N-terminal peptide by cleavage immediately upstream of the inserted FLAG tag (DYKDDDDK), which was positioned between residues S46 and E47.

Peptides were desalted using C18 StageTips (Empore) and dried under vacuum. Peptide samples were analyzed by LC-MS/MS using an UltiMate 3000 RSLC nano system coupled to an Orbitrap Fusion Lumos Tribrid mass spectrometer (Thermo Fisher Scientific). Peptides were loaded onto a C18 trap column (Acclaim PepMap, 75μm × 2 cm, 3μm, 100 Å) and separated on a C18 analytical column (Acclaim PepMap, 50 μm × 15 cm, 2μm, 100 Å) using a linear gradient of 3-45% buffer B over 55min, followed by 45–98% buffer B over 5 min and a 5 min hold at 98% buffer B, at a flow rate of 300nL/min (buffer A: 0.1% formic acid in water; buffer B: 0.1% formic acid in acetonitrile containing 20% water). Survey scans were acquired in positive ion mode over an m/z range of 350–1,800 at a resolution of 60,000 (at m/z 200). Data-dependent acquisition was performed with a 3s cycle time. Precursor ions were fragmented by higher-energy collisional dissociation with 30% normalized collision energy, and fragment ions were detected at a resolution of 15,000. Dynamic exclusion was enabled for 60s with a 10-ppm mass tolerance. Raw data were processed using MaxQuant (version 2.6.4.0) with peptide identification performed by the Andromeda search engine against the Caenorhabditis elegans UniProt reference proteome (UniProt ID: UP000001940), combined with the MaxQuant contaminant database. Label-free quantification (LFQ) was enabled, and a false discovery rate of 1% was applied at both the peptide and protein levels. Peptide MS1 signal intensities were quantified using Skyline (version 25.1).

### Bioinformatic prediction of APP-1/XPNPEP1– and DPF-3/DPP8/9–co-regulated substrates in *C. elegans* and humans

Custom Python scripts were used to predict protein substrates cooperatively regulated by APP-1/XPNPEP1 and DPF-3/DPP8/9. For each protein, three metrics were calculated: the maximum N-terminal cleavability achieved by APP-1/XPNPEP1 alone, by DPF-3/DPP8/9 alone, and by the combined action of both enzymes. Proteins predicted to undergo more extensive processing when both enzymes were present than with either enzyme alone were retained as candidate co-regulated substrates. Functional enrichment analysis was performed using the DAVID platform, and Biological Process enrichment results were exported and visualized as bubble plots in R. The analysis scripts are available upon request.

## Data availability

The mass spectrometry proteomics data have been deposited to the ProteomeXchange Consortium via the PRIDE partner repository, with dataset identifiers PXD072457. RNA-seq datasets have been deposited in the NCBI Sequence Read Archive (SRA) under the BioProject accession number PRJNA1392415.

## Statistics

No statistical method was used to predetermine the sample size. No data were excluded from the analyses. The experiments were not randomized. Investigators were not blinded to group allocation during experiments and outcome assessment. Two-tailed Student’s t-test and one-way ANOVA were used to derive P values for all figures. Data are presented as mean ± s.d., unless otherwise indicated.

## Acknowledgment

We thank members of the Wan laboratory for discussions and Rene Ketting and colleagues (Institute of Molecular Biology, Johannes Gutenberg University, Mainz) for sharing unpublished results. We are also grateful to Jianfeng Li, Fengzhu Wang and colleagues at Sun Yat-sen University for providing yeast two-hybrid plasmids and protocols. This work was supported by the National Natural Science Foundation of China (32570824 and 32370729), the Shenzhen Medical Research Fund (B2302029), the Natural Science Foundation of Guangdong Province (2024A1515012650) to G.W., and the Guangdong Science and Technology Department (2023B1212060028).

## Contributions

B.D., X.N. and G.W. designed the experiments. B.D. and X.N. performed most of the experiments with the help from H.M., C.Z., W.C., S.H., Y. L., K.C., Y.L., Y.Q., H.J. L.S. and X. L. performed the bioinformatic analysis of small RNA-seq data. All authors analyzed the data. G.W. conceived and supervised the project, interpreted the results, and wrote the paper.

## Supplemental Figures and Legends

**Supplemental Figure 1.**
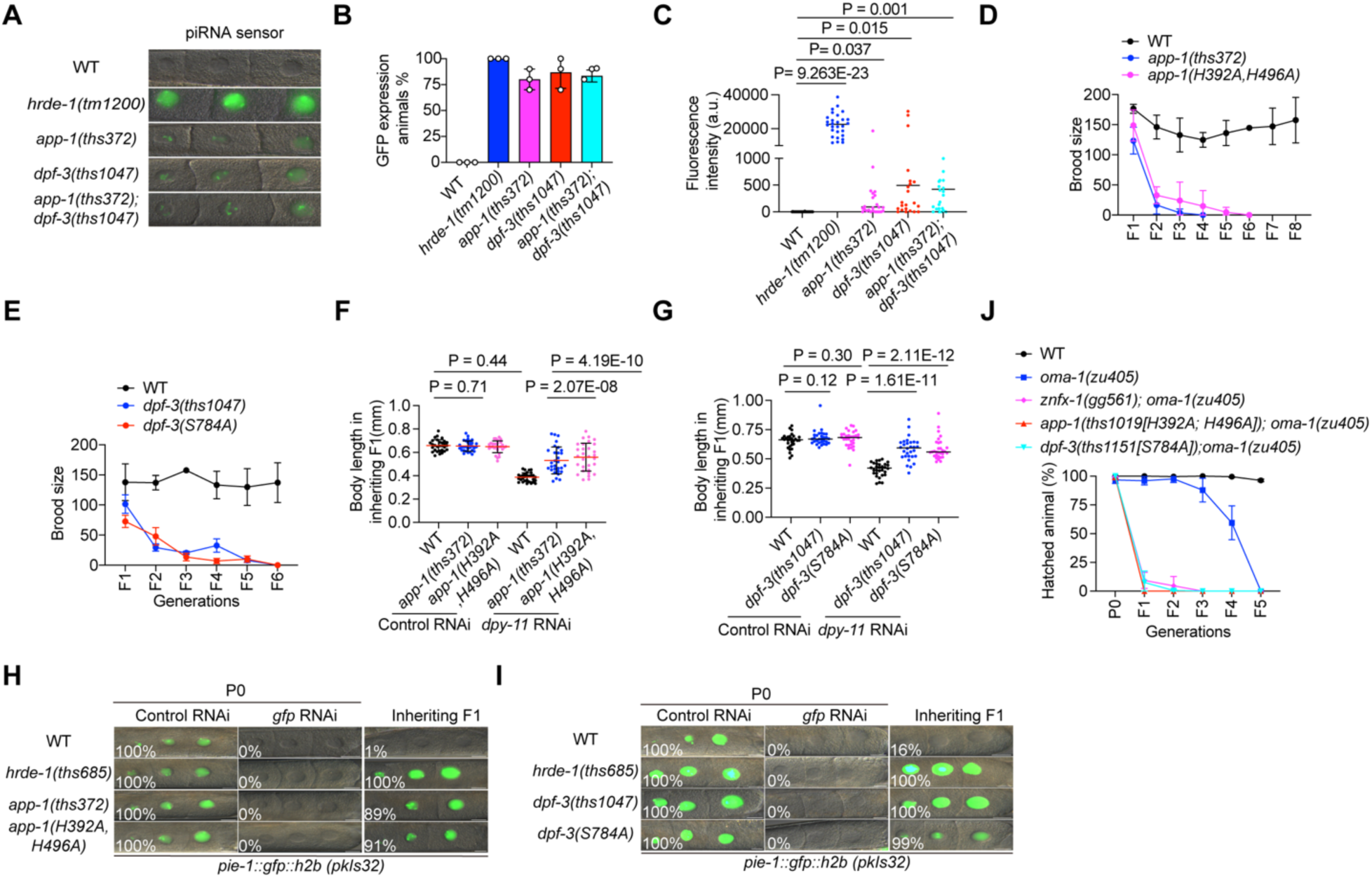
Enzymatic activity of APP-1 and DPF-3 promote small RNA-based epigenetic inheritance. (A) Fluorescence imaging of a piRNA sensor (*mex-5p::GFP::his-58::21UR-1 target::tbb-2 3’UTR II*) in WT, *app-1(-)*, *dpf-3(-)*, and *app-1(-); dpf-3(-)* background. (B-C) Quantification of percentage of animals expressing GFP (B) and fluorescence intensity of GFP (C) with Image J. For panel B, n = 3 biological replicates, with 10 animals scored per replicate. For panel C, n = 30 animals from three independent plates. Student’s t test, two-tailed. (D-E) Animals with indicated genotype were grown at 25°C, brood size was scored for indicated generations. n = 3 biological replicates. Student’s t test, two-tailed. (F-G) Animals with indicated genotype were subjected to *dpy-11* RNAi inheritance assay. The length of L4 stage animals in inheriting F1 were measured with ImageJ. n = 30 animals for each genotype. Student’s t test, two-tailed. (H-I) Animals with indicated genotype expressing *gfp::h2b* in the germline were exposed to *gfp* RNAi, progeny were maintained at no RNAi plates (OP50). The percentage of animals expressing GFP in −1 to −3 oocytes were scored. n = 3 biological replicates (50 animals were scored for each replicate). Scale bar, 10μm. (J) Animals carrying *oma-1(zu405)* were treated with *oma-1* dsRNA and maintained at 20°C. The percentage of hatched progeny were scored across generations. n = 3 biological replicates.

**Supplemental Figure 2.**
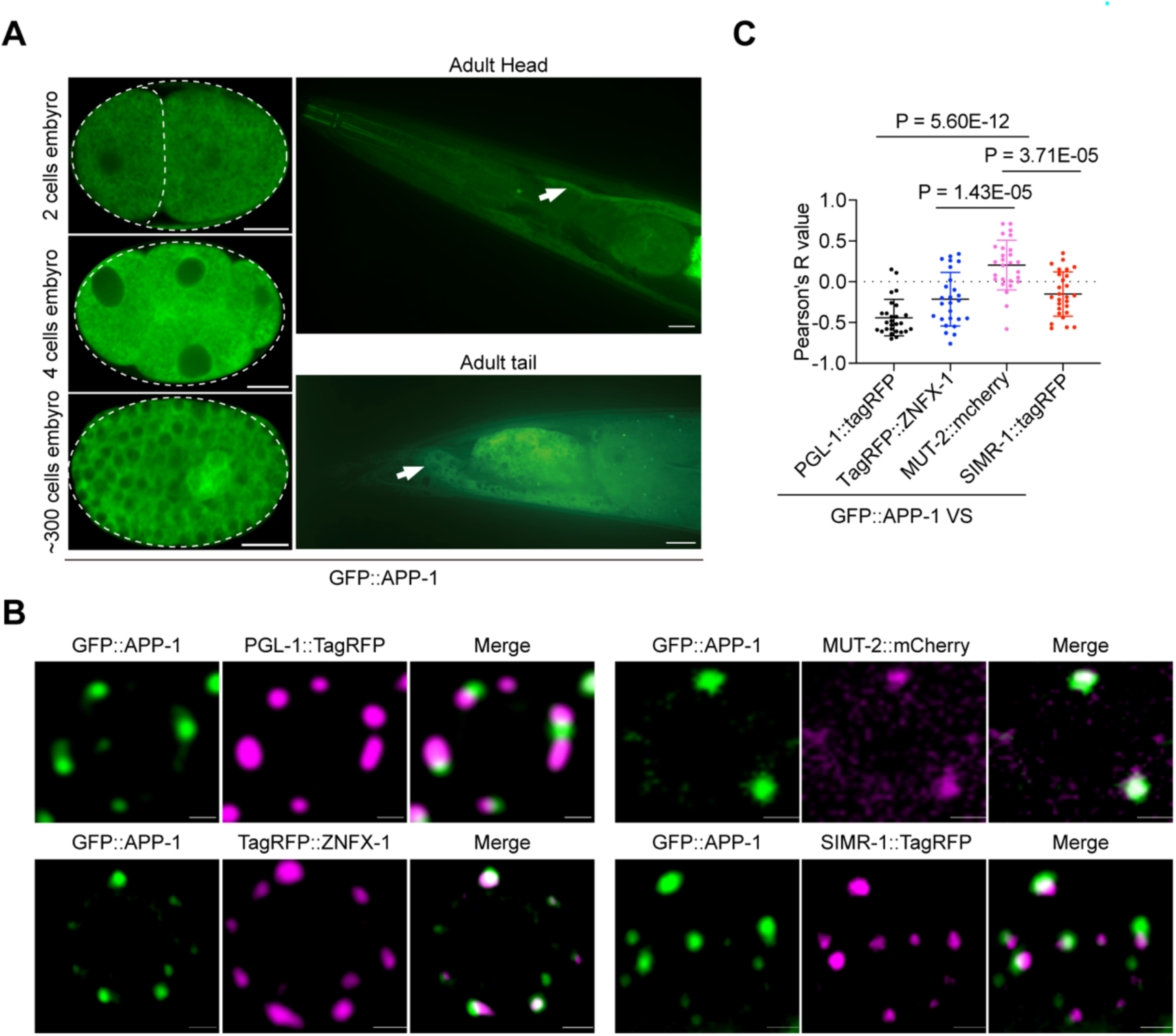
The localization of APP-1. (A) Fluorescence imaging of embryos or young adult animals expressing GFP::APP-1. n > 3 animals. Scale bar, 10μm. (B) Fluorescence imaging of pachytene region of young adult germline expressing indicated proteins. Scale bar, 1μm. (C) Image J analysis to measure the degree of colocalization of indicated proteins. n = 27 (3 granules per cell, 3 cells per animals, 3 animals per sample). Student’s t test, two-tailed.

**Supplemental Figure 3.**
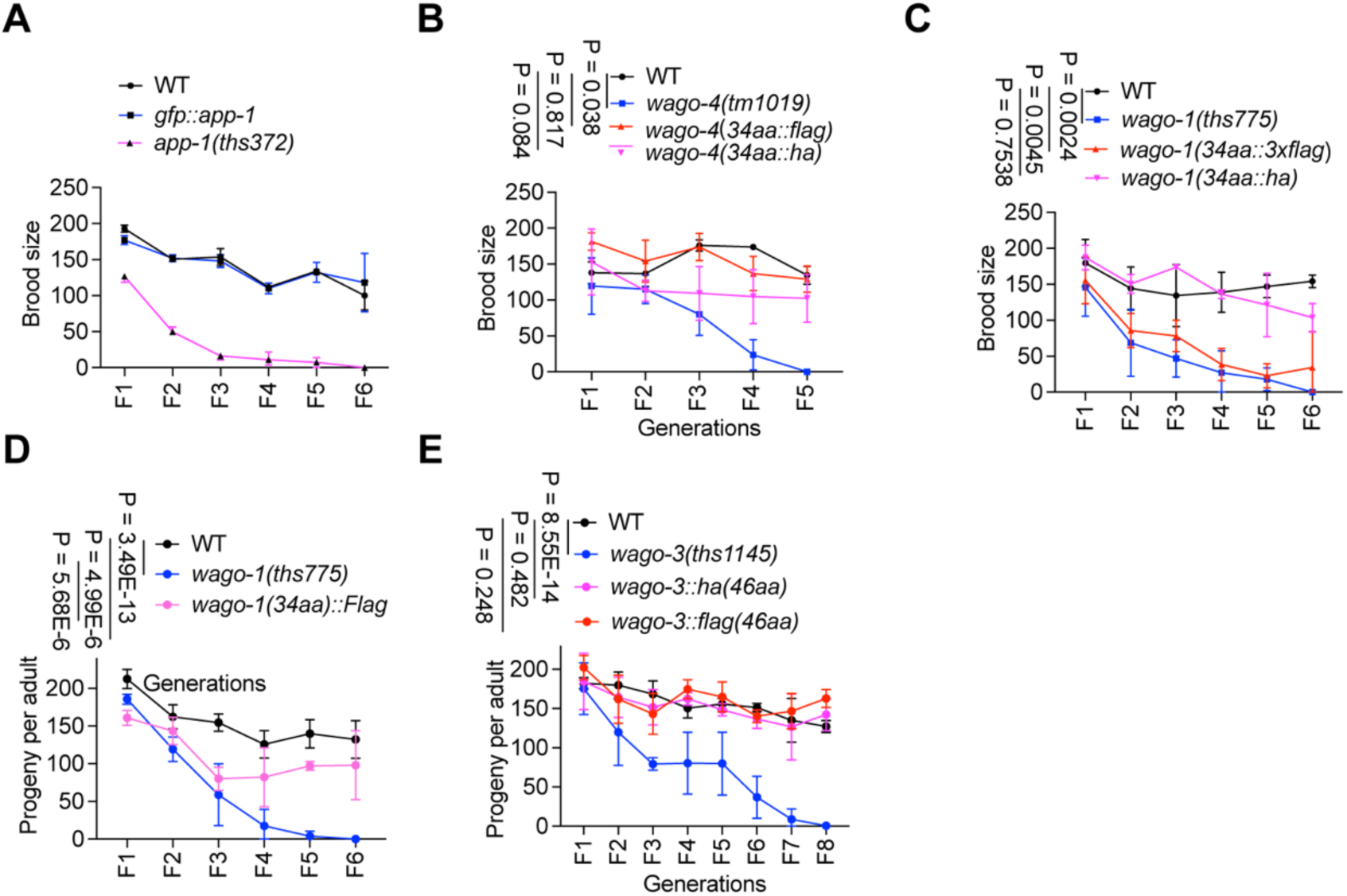
Introduction of GFP, FLAG, and HA tags into endogenous gene loci using CRISPR/Cas9 generated functional fusion proteins. (A-E) Animals with indicated genotype were grown at 25 °C, brood size was scored for indicated generations. n = 3 biological replicates. P values were determined by one-way ANOVA.

**Supplemental Figure 4.**
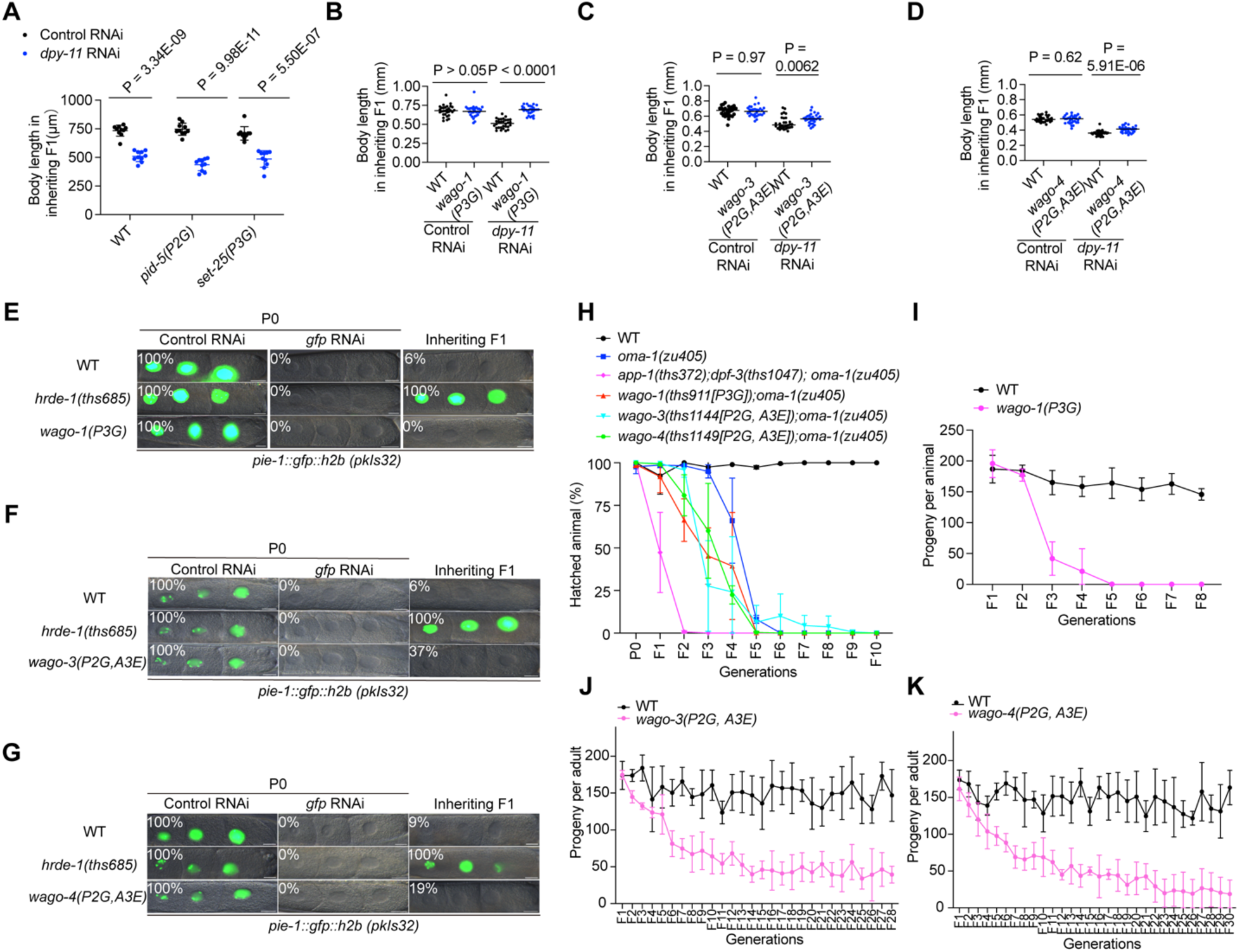
Identification of WAGO-1, WAGO-3, and WAGO-4 as potential substrates of APP-1 and DPF-3. (A-D) WT animals and animals expressing predicted APP-1 and DPF-3 subtrates with processing defect mutations were treated with *dpy-11* RNAi, the progeny were grown on control RNAi (OP50) plates. Body length was measured for the progeny at L4 stage. n = 3 biological replicates (30 animals were measured per genotype). Student’s t test, two tailed. (E-G) WT, *hrde-1(-)*, and *wago-1(P3G)* (E), or WT, *hrde-1(-)*, and *wago-3(P2G, A3E)* (F), or WT, *hrde-1(-)*, and *wago-4(P2G, A3E)* (G) animals expressing *gfp::h2b* (*pkIS32*) in the germline were treated with *gfp* RNAi, progeny were grown on non-RNAi plates. The percentage of GFP expression animals in P0 and inheriting F1 generations were scored under a 10X subject in a Leica SP8 inverted microscrope. n = 3 biological replicates ( For each replicate, 50 animals were scored). Scale bar, 10μm. (H) Animals with *oma-1(zu405)* in the background were subjected to *oma-1* RNAi and maintained at 20°C, the precentage of hatched animals were scored for indicated generations. n = 3 biological replicates. (I-K) Animals were grown at 25°C, brood size were scored for each generation. n = 3 biological replicates.

**Supplemental Figure 5.**
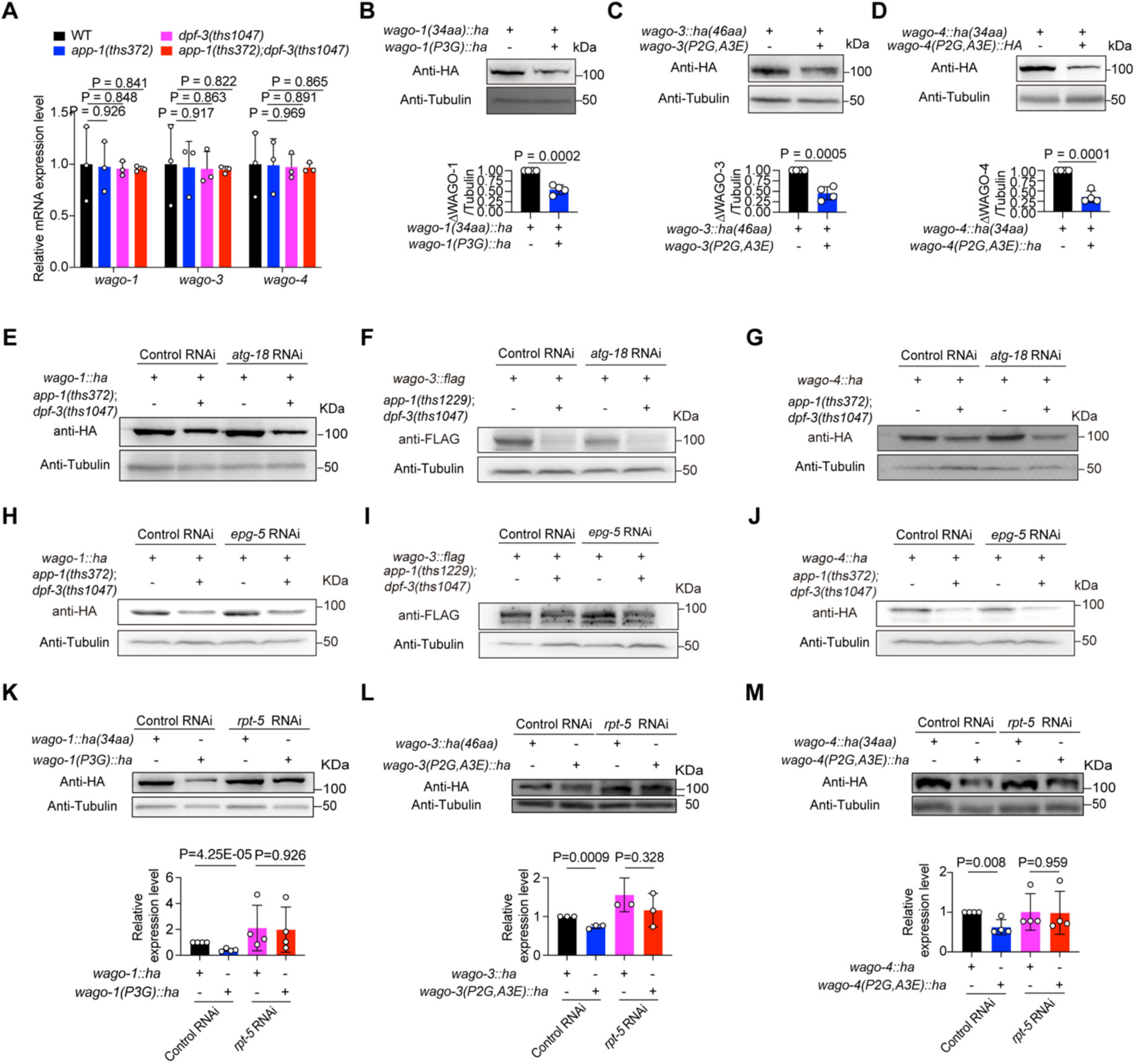
Loss of predicted APP-1 and DPF-3 processing leads to proteasome-mediated, but not autophagy-mediated, degradation of WAGO-1/3/4. (A) qRT-PCR analysis of *wago-1*, *wago-3*, and *wago-4* mRNA level in *app-1(-)*, *dpf-3(-)*, *app-1(-)*; *dpf-3(-)* animals. n = 3 biological replicates. Student’s t test, two tailed. (B-D) Western blots detecting WAGO-1, WAGO-3, and WAGO-4 protein levels in WT and processing-defective mutants (top). Quantification of WAGO-1/3/4 protein levels normalized to Tubulin is shown below. n = 4 biological replicates. (E-G) Western blots detecting WAGO-1, WAGO-3, and WAGO-4 in WT and *app-1(-)*; *dpf-3(-)* animals treated with control RNAi or *atg-18* RNAi. n = 2 biological replicates. (H-J) Western blots detecting WAGO-1, WAGO-3, and WAGO-4 in WT and *app-1(-)*; *dpf-3(-)* animals treated with control RNAi or *epg-5* RNAi. n = 2 biological replicates. (K–M) Western blots detecting WAGO-1, WAGO-3, and WAGO-4 in WT and processing-defective mutants treated with control RNAi or *rpt-5* RNAi (top). Quantification of WAGO-1/3/4 protein levels normalized to Tubulin is shown below. n = 3 biological replicates. Student’s t test, two tailed.

**Supplemental Figure 6.**
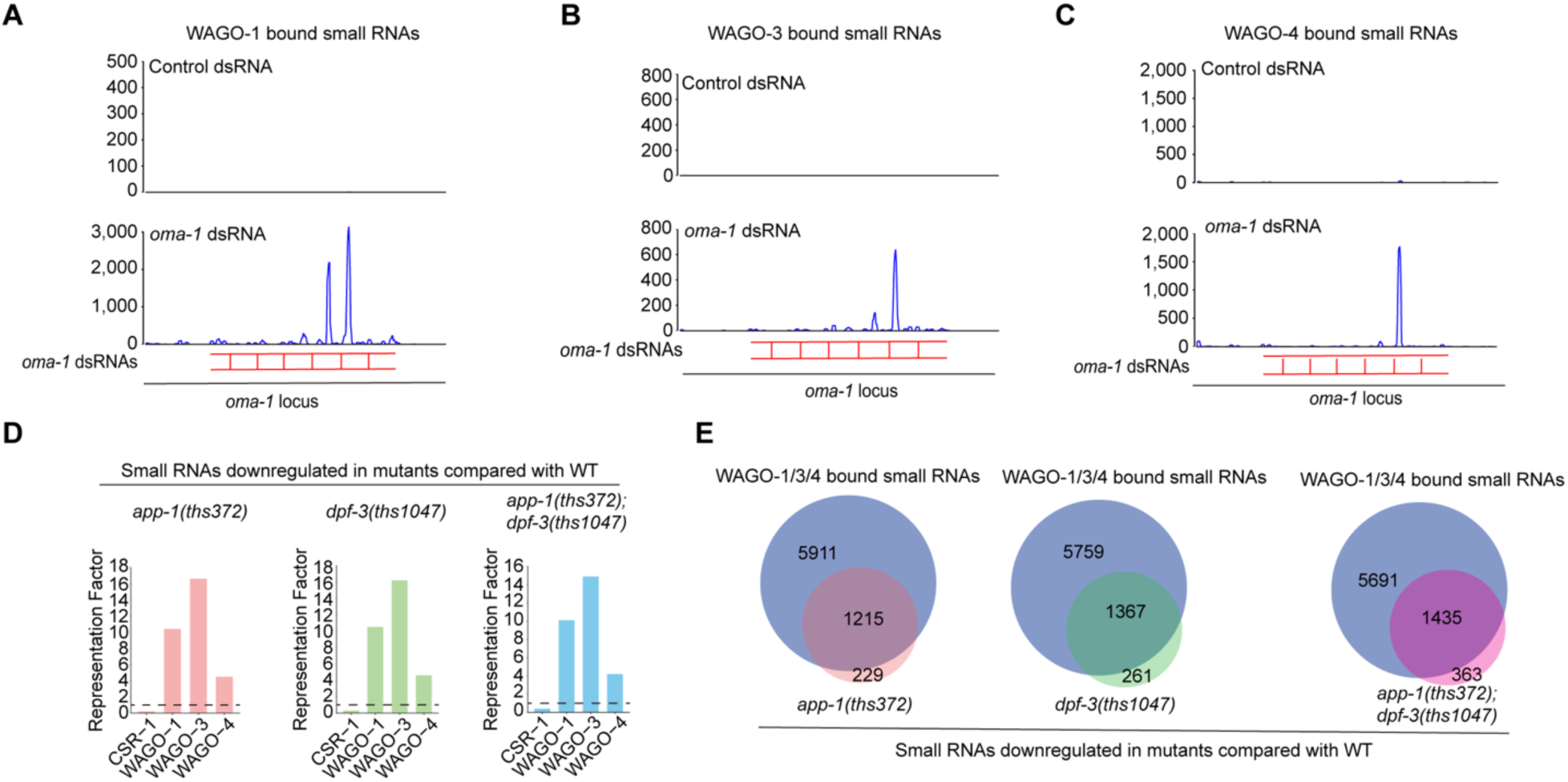
Endogenous siRNAs reduced upon APP-1/DPF-3 loss are predominantly WAGO-1/3/4-bound siRNAs. (A-C) Animals expressing FLAG::WAGO-1, FLAG::WAGO-3, or FLAG::WAGO-4 were treated with control dsRNA or *oma-1* dsRNA. Associated *oma-1* siRNAs bound to FLAG::WAGO-1 (A), FLAG::WAGO-3 (B), and FLAG::WAGO-4 (C) were quantified by small RNA-seq. n = 2 biological replicates. (D) Representative factor analysis comparing genes targeted by WAGO-1/3/4-associated endogenous siRNAs with genes showing reduced endogenous siRNAs in *app-1(-)*, *dpf-3(-)*, and *app-1(-)*; *dpf-3(-)* animals. (E)Venn diagram showing the overlap between genes targeted by WAGO-1/3/4-associated endogenous siRNAs and genes with reduced endogenous siRNAs in *app-1(-)*, *dpf-3(-)*, and *app-1(-)*; *dpf-3(-)* animals.

**Supplemental Figure 7.**
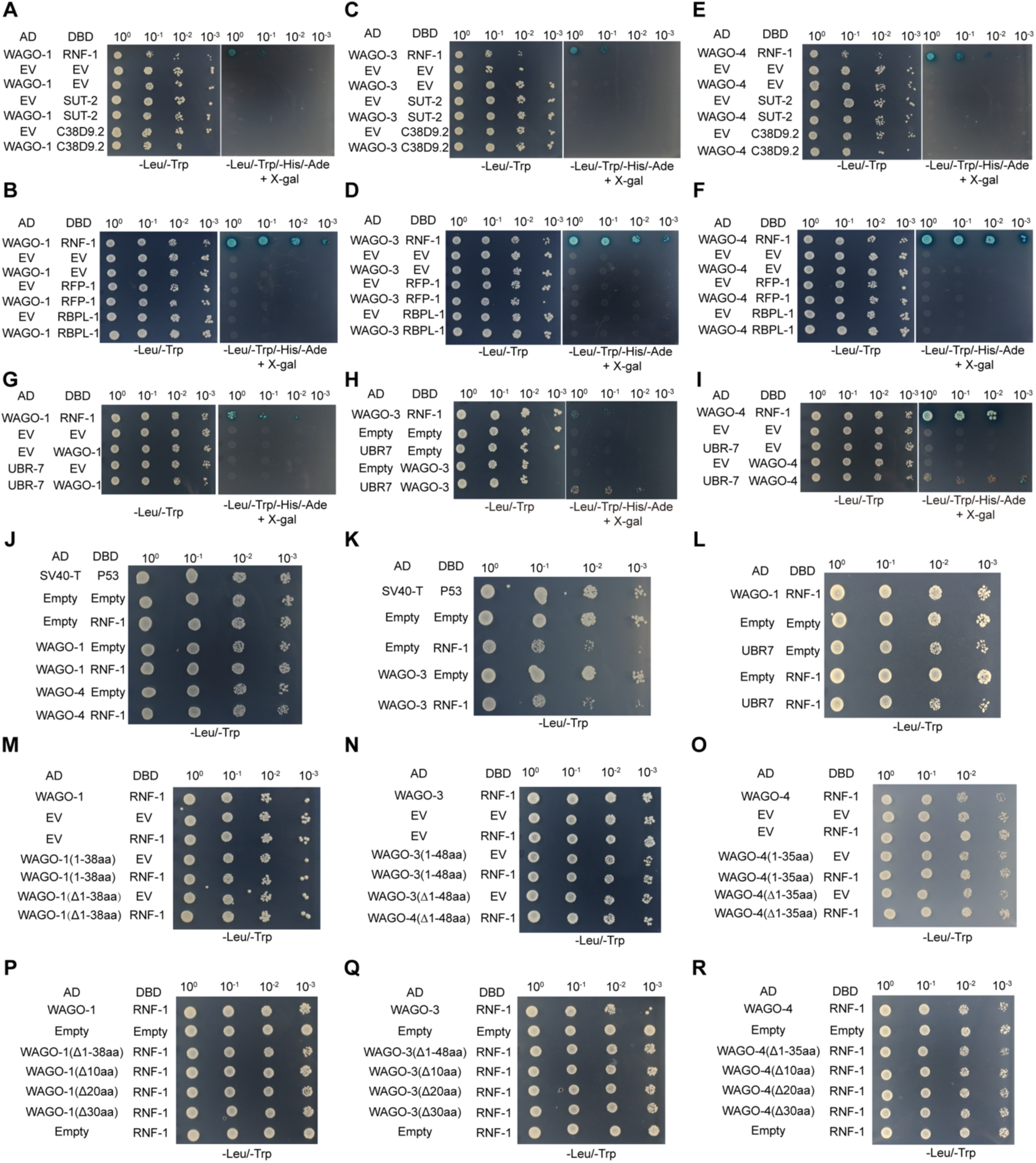
Yeast two-hybrid assays testing interactions between WAGO-1/3/4 and five candidate E3 ligases. (A-I) Yeast two-hybrid assays testing the interactions between WAGO-1/3/4 and five candidate E3 ligases. Note, Images in panels A-F were acquired 3 days after yeast were transferred to SD/−Leu/−Trp/−Ade/−His/X-gal medium. Images in panels G-I were acquired 6 days after transfer to the same medium. (J-K) Yeast transformed with plasmids expressing the indicated proteins and grown on - Leu/-Trp plates to confirm comparable growth, related to Figure 5B-C. (G-I) Yeast transformed with plasmids expressing the indicated proteins and grown on - Leu/-Trp plates to confirm comparable growth, related to Figure 5M. (M-O) Yeast transformed with plasmids expressing the indicated proteins and grown on-Leu/-Trp plates to confirm comparable growth, related to Figure 5N-P. (P-R) Yeast transformed with plasmids expressing the indicated proteins and grown on - Leu/-Trp plates to confirm comparable growth, related to Figure 5Q-S.

**Supplemental Figure 8.**
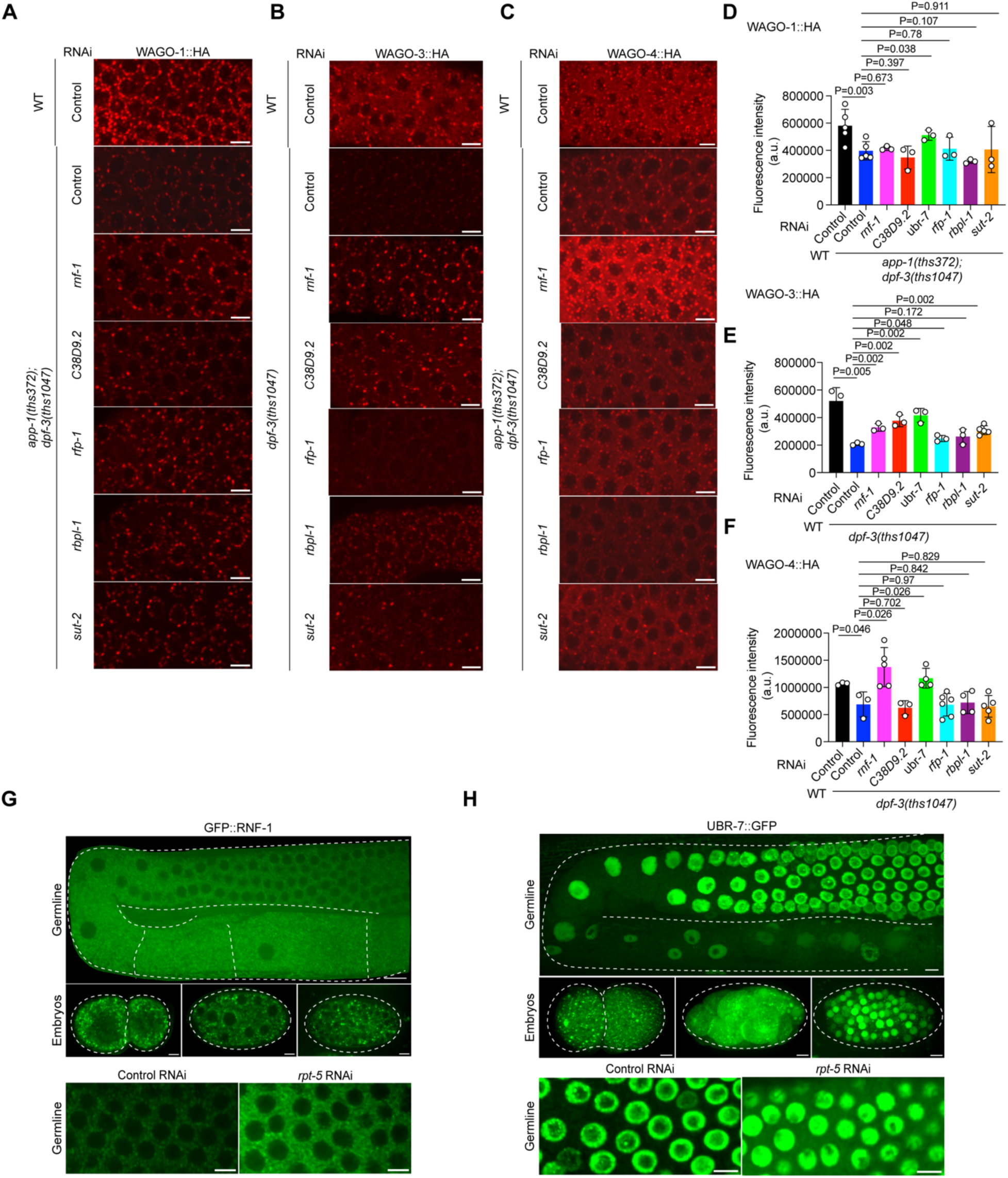
Identification of E3 ligases that regulate the stability of unprocessed WAGO-1, WAGO-3, and WAGO-4. (A-C) Immunofluorescence analysis of WAGO-1, WAGO-3, and WAGO-4 in WT, *app-1(-)*; *dpf-3(-)*, or *dpf-3(-)* animals treated with control RNAi or RNAi against candidate E3 ligases. Scale bar, 5μm. (D-F) Quantification of WAGO-1, WAGO-3, and WAGO-4 fluorescence intensities shown in (A-C) and Figure 5D-F. (G) Localization of GFP::RNF-1, with WT animals imaged under identical exposure conditions as controls. n > 3 animals. Scale bar, 5μm. (H) Localization of GFP::UBR-7, with WT animals imaged under identical exposure conditions as controls. n > 3 animals. Scale bar, 5μm.

**Supplemental Figure 9.**
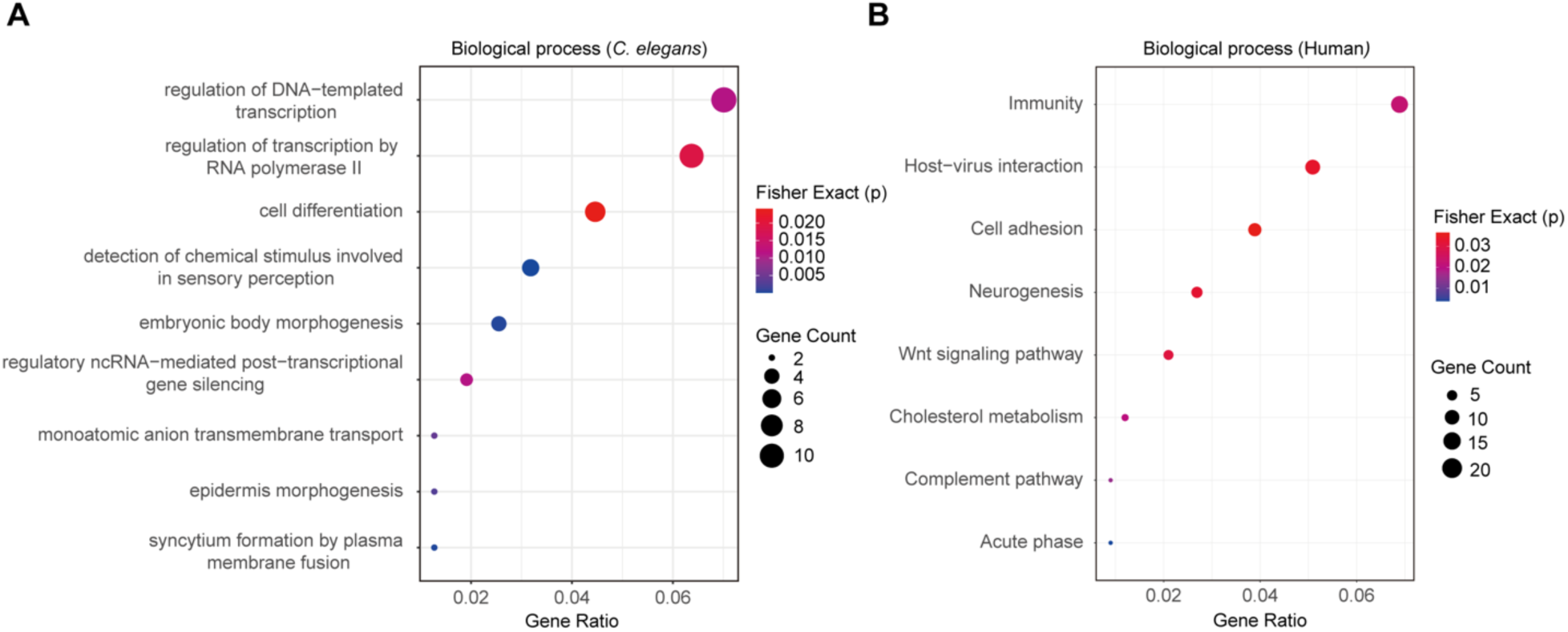
Biological process enrichement analysis of predicted APP-1 and DPF-3 cooperatively processed substrates in *C. elegans* and human.

